# The 3D Structural Architecture of the Human Hand Area is Non-Topographic

**DOI:** 10.1101/2022.07.28.501903

**Authors:** Juliane Doehler, Alicia Northall, Peng Liu, Alessio Fracasso, Anastasia Chrysidou, Oliver Speck, Gabriele Lohmann, Thomas Wolbers, Esther Kuehn

## Abstract

The functional topography of the human primary somatosensory cortex (S1) hand area is a widely studied model system to understand sensory organization and plasticity. It is so far unclear whether or not the underlying 3D structural architecture also shows a topographic organization. We used 7T MRI data to quantify layer-specific myelin, iron and mineralization in relation to population receptive field maps of individual finger representations in Brodman area 3b (BA 3b) of human S1 in female and male younger adults. This 3D description allowed us to identify a characteristic profile of layer-specific myelin and iron deposition in the BA 3b hand area, but revealed an absence of structural differences, an absence of low-myelin borders, and high similarity of 3D microstructure profiles between individual fingers. However, structural differences and borders were detected between the hand and face areas. We conclude that the 3D structural architecture of the human hand area is non-topographic, unlike in some monkey species, which suggests a high degree of flexibility for functional finger organization and a new perspective on human topographic plasticity.

**Significance Statement:** Using ultra-high field MRI, we provide the first comprehensive *in vivo* description of the 3D structural architecture of the human BA 3b hand area in relation to functional population receptive field (pRF) maps. High similarity of precise finger-specific 3D profiles, together with an absence of structural differences and an absence of low-myelin borders between individual fingers, reveal the 3D structural architecture of the human hand area to be non-topographic. This suggests reduced structural limitations to cortical plasticity and reorganization, and allows for shared representational features across fingers.

## Introduction

In the mammalian brain, the topographic architecture of the primary somatosensory cortex (S1) is often studied as a model system to understand cortical functional organization and plasticity (e.g. Florence et al., 1997; Feldman and Brecht, 2005; Kuehn and Pleger, 2020). Due to their clear and fine-grained architectures, the most studied topographic areas within S1 are whisker representations in rodents (Petersen, 2007; Feldman and Brecht, 2005), and finger representations in monkeys and humans (Shoham and Grinvald, 2001; Blake et al., 2002; Kuehn et al., 2018a; Pleger et al., 2016). While the microstructural features of these areas in relation to functional map architectures have been widely studied in rodents and monkeys (e.g. Meyer et al., 2013; Welker, 1976; Welker and Woolsey, 1976; Peters, 2009, Qi and Kaas, 2004; Jain, Catania and Kaas, 1998), this remains undescribed in humans. This knowledge gap prevents us from gaining a full understanding of the neuronal mechanisms that underlie topographic organization and plasticity in humans.

Physiological studies show that low-myelin borders (so-called septa) separate major body part representations (Manger et al., 1997) and finger representations in Brodmann area 3b (BA 3b, homologue of rodent S1) of macaque monkeys (Qi and Kaas, 2004) and owl monkeys (Jain, Catania and Kaas, 1998), as well as whisker representations in rodents (Woolsey and Van der Loos, 1970). The role of such septa is complex and ranges from functional separation to sensory encoding; it has also been suggested that septa may limit cortical plasticity (Sereno, 2005). Recently, low-myelin borders have been identified in human BA 3b between major body part representations (Glasser et al., 2016; Kuehn et al., 2017). However, it is so far unknown whether low-myelin borders also separate functional single finger representations in human BA 3b, or whether the human hand area can be regarded as one, structurally homogeneous cortical field (Sereno et al., 2022), where (i) single finger septa and (ii) structural gradients (previously related to large-scale functional organization, e.g. Huntenburg et al., 2018; Paquola et al., 2019; Valk et al., 2022; Wagstyl et al., 2015) are absent, and (iii) single finger representations appear structurally similar.

Conceptually, this question is important because it touches on a fundamental aspect of brain organization. In the somatosensory system, unlike in the visual system, topographic representations are discontinuous. That is, single finger representations (e.g. Schweizer et al., 2008; Martuzzi et al., 2014; Schellekens et al., 2021) and major body part representations (Sereno et al., 2022), such as the hand and the face, are distinctly represented as has been shown with fMRI. This representation of body parts reflects the discontinuous shape of the body in the real world, where the hand and the face, for example, are spatially distant (and can move independently) even though they cover neighboring areas in S1. Therefore, it has been suggested that low-myelin borders in BA 3b separate representations that are nearby in the cortex but distant in the real world (Kuehn et al., 2017), such as the hand and face. Alternatively, low-myelin borders may separate representations that are nearby in the cortex and nearby in the real world, but that sometimes receive distinct sensory inputs, such as individual fingers. Answering this question will facilitate relating cortical myeloarchitectonic features to real world features.

We here used parcellation-inspired techniques to characterize finger-specific structural features of the hand representation in human BA 3b, not only along the surface (two dimensions) but also in depth (three dimensions). Quantitative MRI proxies were used to estimate layer-specific myelin, diamagnetic and paramagnetic tissue substances (such as calcium and iron) and overall mineralization. Participants underwent a tactile stimulation protocol to obtain population receptive field (pRF) characteristics to investigate whether the 3D structural architecture of the human hand area is homogenous, or whether there are systematic microstructural differences between BA 3b finger representations. Targeting these questions provides us with critical information on topographic stability and/or plasticity in conditions of health and disease.

## Methods

### Participants

24 healthy volunteers underwent 7T MRI. Due to severe head motion during scanning, 4 participants were excluded from the study, leaving a total of 20 participants for analysis (10 females, mean age = 25 +/- 3 years). The participant number was motivated by previous layer-dependent 7T MRI studies using quantitative in-vivo proxies to describe the structural architecture of human BA 3b (Dinse et al., 2015, Kuehn et al., 2017), and is well above previously reported sample sizes. According to the Edinburgh handedness questionnaire (Oldfield, 1971), all participants were right-handed (laterality index ranging from +33 to +100, M = 82 +/- 21 SD). Chronic illness, central acting medications and MRI contraindications (e.g., active implants, non-removable metallic objects, claustrophobia, tinnitus or hearing impairments, consumption of alcohol/drugs) were a priori exclusion criteria.

Participants showed no anomalies of sensory perception (e.g., numbness, tingling sensations) or motor movement (e.g., reduced motor control, restricted finger movement), and no diagnoses of diabetes or hypertension. No professional musicians participated, given evidence of enlarged cortical hand representations and superior tactile perception in string and piano players (Elbert et al., 1995; Ragert et al., 2004; Schwenkreis et al., 2007). Finally, none of the participants showed signs of cognitive impairments as indicated by the ’Montreal Cognitive Assessment’ (Nasreddine et al., 2005; M = 29 +/- 1 SD, scores ranging from 26 to 30). Participants were recruited from the database of the DZNE Magdeburg, Germany. All participants gave their written informed consent and were paid for their participation. The study was approved by the Ethics committee of the Otto-von-Guericke University Magdeburg. fMRI data were partly published in a previous study (Liu et al., 2021).

### General Procedure

All participants took part in two appointments: (i) structural MRI session and (ii) functional MRI session (see **Figure 1** for experimental design and analysis pipeline of MRI data).

**Figure 1.**
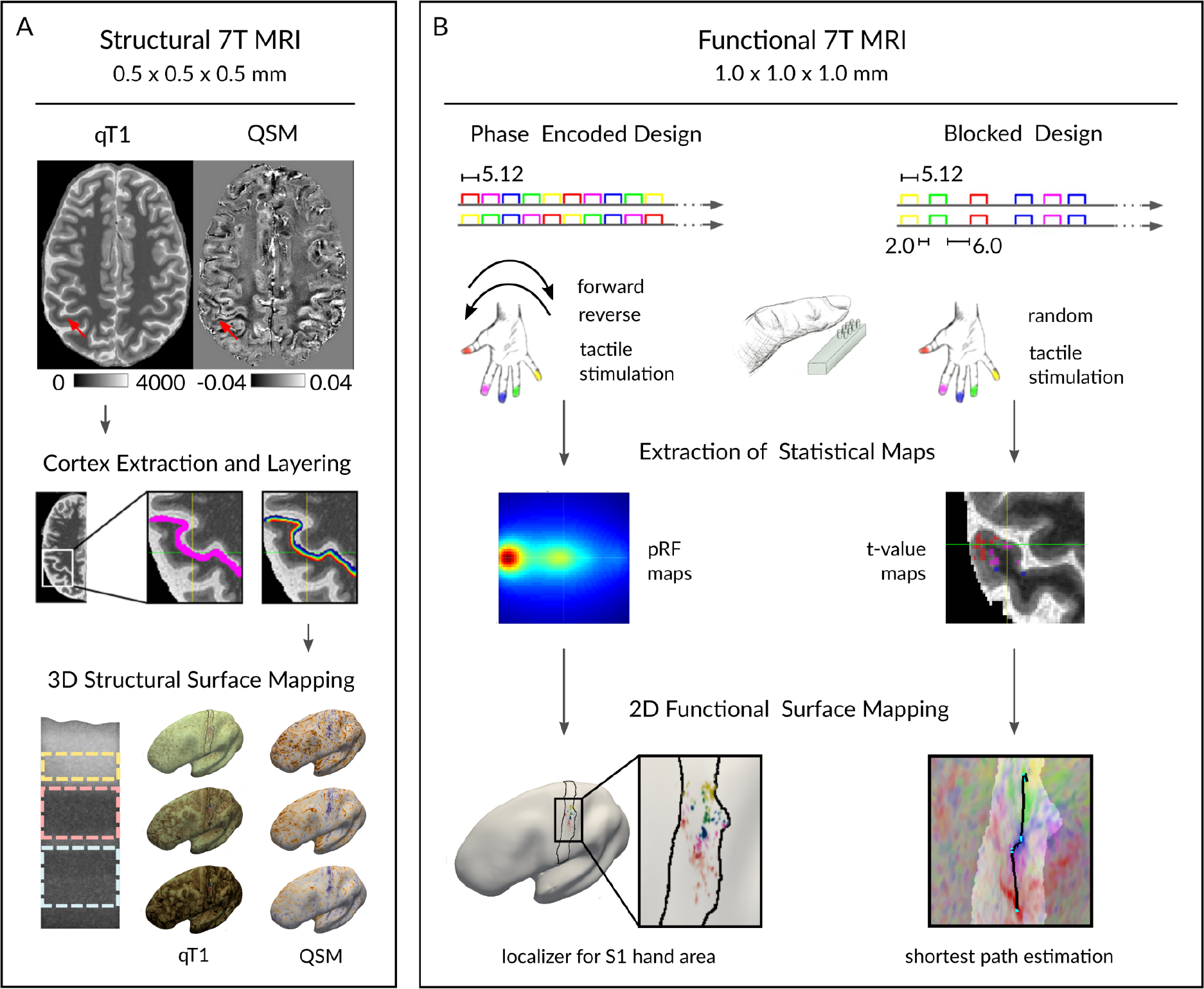
Overview Experimental Design. A total of n=20 participants took part in structural and functional 7T MRI data acquisition. (A) MP2RAGE and susceptibility-weighted imaging sequences of participants were acquired to estimate high-resolution quantitative T1 maps (qT1) and quantitative susceptibility maps (QSM), respectively. Red arrows indicate the region of interest (left hand representation in BA 3b). The MP2RAGE sequence was used to extract the cortex (magenta color), and to layer the cortex into 21 cortical depths (rainbow color) where qT1 and QSM values were sampled to derive cortical depth-dependent profiles. These profiles were averaged into three anatomically-relevant compartments (creme: outer, light pink: middle, light blue: inner) based on *ex vivo in vivo* validated data (see details below, Dinse et al., 2015). Extracted qT1 and QSM values were mapped onto the individual’s inflated cortical surface. Myelin staining was remodeled according to Dinse et al. (2015). (B) All participants underwent tactile stimulation during scanning using both a phase-encoded design and a blocked design. Five independently-controlled MR-compatible piezoelectric stimulators were used to stimulate the five fingertips of the right hand. Phase encoded data were used to calculate population receptive field (pRF) properties, which were used to localize the finger representations in BA 3b. Independent blocked design data were used to calculate t-maps for shortest path estimation.

### MRI Assessment

#### MR Sequences

MRI data were acquired at a 7-Tesla MAGNETOM scanner (Siemens Healthcare, Erlangen, Germany) equipped with a 32 Channel Nova Medical head coil. First, MP2RAGE (Marques et al., 2010) whole brain images were acquired at 0.7 mm isotropic resolution (240 sagittal slices, field-of view read = 224 mm, repetition time = 4800 ms, echo time =2.01 ms, inversion time TI1/TI2 = 900/2750 ms, flip angle = 5◦/3◦ , bandwidth = 250 Hz/Px, GRAPPA 2), and MP2RAGE part brain images (covering the sensorimotor cortex) were acquired subsequently at 0.5 mm isotropic resolution (208 transversal slices, field-of-view read = 224 mm, repetition time = 4800 ms, echo time = 2.62 ms, inversion time TI1/TI2 = 900/2750 ms, flip angle = 5◦/3◦, bandwidth = 250 Hz/Px, GRAPPA 2, phase oversampling = 0%, slice oversampling = 7.7%). Then, we acquired uncombined susceptibility-weighted imaging (SWI, Haacke at al., 2004) data with part brain coverage of sensorimotor cortex at 0.5 mm isotropic resolution using a 3D gradient-echo pulse sequence (208 transversal slices, field-of-view read = 192 mm, repetition time = 22 ms, echo time = 9.00 ms, flip angle = 10◦, bandwidth = 160 Hz/Px, GRAPPA 2, phase oversampling = 0%, slice oversampling = 7.7%). The total scanning time was approximately 60 minutes. Functional data were collected in a separate scanning session. First, shimming was performed, before two EPIs with opposite phase-encoding (PE) polarity were acquired. Functional EPI sequences (gradient-echo) were acquired using the following parameters: voxel resolution of 1 mm isotropic, field-of-view read: 192 mm, repetition time = 2000 ms, echo time = 22 ms, GRAPPA 4, interleaved acquisition, 36 slices.

#### fMRI Task

Five independently-controlled MR-compatible piezoelectric stimulators (Quaerosys, http://www.quaerosys.com) were used to stimulate the five fingertips of the right hand of the participants in the scanner (Schweisfurth et al., 2015, 2014, 2011). A stimulator was attached to each fingertip of the right hand, using a custom-built, metal-free applicator to adjust to individual hand and finger sizes. Each stimulator consisted of 8 individually-controlled pins arranged in a 2x4 matrix, covering 2.5x9 mm^2^ of skin (see **Figure 1**). Vibrotactile stimulation was applied at a frequency of 16 Hz (Schweizer et al., 2008), and stimulation intensity was adjusted individually for each participant and each finger to 2.5 times the individual tactile detection thresholds. To minimize adaptation-related variation in map activity between participants, two randomly selected pins were raised once at a time, yielding 16 pin combinations per second (Schweisfurth et al., 2015, 2014, 2011).

Participants first underwent two phase-encoded protocols (runs 1 and 2 of the experiment), which included 2 runs of 20 cycles each. Each cycle lasted 25.6 seconds where each fingertip was stimulated 20 times for 5.12 seconds. Stimulation was applied either in a forward (D1->D5) or in a reverse order (D5->D1, see **Figure 1**). Half of the participants started with the forward-run, while the other half started with the reverse-run. One run comprised 256 scans (512 seconds for a TR of 2 seconds), and lasted for 8 minutes and 31 seconds. Participants were instructed to covertly count short, randomly distributed pauses during the tactile stimulation (duration 180 ms). We used the same number of gaps per finger, resulting in 15 gaps in total per run. Participants then underwent the blocked-design protocol (runs 3 and 4 of the experiment), which included 6 conditions: Stimulation to D1, D2, D3, D4, D5, and a rest condition with no stimulation. The same task instructions and stimulation protocol as described for the phase-encoded paradigm was used, although fingers were stimulated in a pseudo-random way, where fingers were not stimulated more than two times in a row. Between two subsequent stimulations, there was a 2 seconds pause (in 70% of the trials), or a 6 seconds pause (in 30% of the trials), which was counterbalanced across fingers. Each finger was stimulated 10 times. One run comprised 208 scans, and lasted for 6.56 minutes. The blocked-design run was repeated twice. Subsequently, two runs were recorded where a one-TR stimulation of all five fingers was followed by a 11-TR resting phase without any stimulation. The sequence was repeated 10 times for both runs. All together, functional measurements took approximately 40 minutes.

### MRI Analyses

#### Structural Data Processing Preprocessing

Structural data quality was evaluated by two independent raters and data showing severe artifacts (i.e., of n=3 participants) were excluded. We only used data showing mild truncation artifacts (not affecting S1), or no artifacts at all. Quantitative susceptibility maps (QSM) were reconstructed from coil-uncombined SWI magnitude and phase images using the Bayesian multi-scale dipole inversion (MSDI) algorithm (Acosta-Cabronero et al., 2018) implemented in the software qsmbox (version v3, freely available for download: https://gitlab.com/acostaj/QSMbox, Acosta-Cabronero et al., 2018) written for Matlab (R2017b, The MathWorks Inc., Natick, MA, 2017). Structural MP2RAGE and QSM images were then processed using JIST (Lucas et al., 2010) and CBS Tools (Bazin et al, 2014) as plug-ins for the research application MIPAV (McAuliffe et al, 2001).

#### Registration

First, the qT1 and QSM slab images were co-registered to the up-sampled (0.7 to 0.5 mm isotropic) whole brain qT1 image. To precisely register the qT1 slab image onto the resampled whole brain qT1 image, we combined linear transformation (MIPAV v7.3.0, McAuliffe et al., 2001, Optimized Automated Registration Algorithm: 6 degrees of freedom, cost function of correlation ratio) and non-linear deformation (ANTs version 1.9.x-Linux, Avants et al., 2011, embedded in cbstools wrapper Embedded Syn, cost function of cross correlation) in one single step, using nearest neighbor interpolation. For registration of the QSM slab image, we applied a combination of rigid and affine automated registration using the software ITK-SNAP (version 3.6.0, freely available for download at www.itksnap.org). The registration quality of resulting qT1 and QSM slab images was evaluated by two independent raters. The generated registration matrix for the qT1 slab image was then applied to the UNI and INV2 images.

#### Segmentation

qT1 slab and whole-brain images were fused using a weighted combination of images, resulting in one whole-brain structural image with improved resolution in the sensorimotor cortex. Using the merged images, brain surrounding tissues (i.e., skull and dura mater) were removed and resulting brain masks were manually refined (using both the qT1 and UNI images) to ensure that all non-brain matter was removed from the region of interest. The cortex was then segmented using the UNI image as input for the TOADS algorithm (Bazin and Pham, 2007) to estimate each voxel’s tissue membership probability. The results were used as input for the CRUISE algorithm (Han et al., 2004) to estimate the inner and outer gray matter (GM) borders (i.e. to the WM and CSF, respectively). The resulting level set images (surfaces in Cartesian space using a level set framework, Sethian, 1999) were optimized to precisely match S1, by thresholding the maximum values of the inner and outer level set images to -2.8 and -0.2, respectively.

#### Layering and Surface Mapping

The level set images were then used to generate individual surfaces for cortical mapping. Intra-cortical qT1 and QSM values, used as proxies for myelin (Stueber, 2014) and mineralization (Acosta-Cabronero et al., 2016; Acosta-Cabronero et al., 2018, Betts et al., 2016), respectively, were estimated in reference to individual cortical folding patterns using the validated equivolume model (Waehnert et al., 2014; Waehnert et al., 2016). The cortical sheath was initially divided into 21 level set surfaces where we sampled qT1 and QSM values to derive cortical depth-dependent profiles. Finally, the extracted values were mapped onto the individual’s inflated cortical surface (method of closest point, Tosun et al., 2014). Please note, for cortical depth-dependent mapping of quantitative values, we used the non-merged high-resolution qT1 and QSM slab data to ensure high data quality. We extracted three different parameter maps from the QSM data (negative QSM (nQSM), positive QSM (pQSM), absolute QSM (aQSM)) to estimate information on different underlying tissue properties (diamagnetic tissue contrast for nQSM, paramagnetic tissue contrast for pQSM, level of mineralization for aQSM).

Extracted cortical depth-dependent profiles were further averaged into fewer compartments following two different approaches: 1) To ensure comparability of our results to previous data, the three most superficial and the two deepest profiles were removed to reduce partial volume effects (Tardif et al., 2015), before the remaining 16 profiles were averaged into four equally-spaced cortical depth compartments (in the following referred to as ‘equally-spaced layers’: superficial, outer-middle, inner-middle, deep). 2) We followed recent *ex vivo in vivo* validation studies in S1 that allow definition of anatomically-relevant cortical compartments from ultra-high resolution MRI data (Dinse et al., 2015, see **Figure 2**). We used previously validated myelin profiles of BA 3b (Dinse et al., 2015) to identify cortical compartments based on averaged qT1 profiles (at the group level) by plotting histological data as well as modeled and *in vivo* MRI data of Dinse et al. (2015) in reference to our *in vivo* MRI data. Calculating minima and maxima of the first derivative of the initial qT1 profile (including 21 compartments) allowed us to extract three data-driven cortical-depth compartments that are anatomically relevant: After removing the two deepest layers (where qT1 stabilized and a plateau was reached, see **Figure 2**), the remaining 19 compartments were averaged into an inner (eight deepest layers), middle (seven layers) and outer (four most superficial layers) cortical-depth compartment, where based on Dinse et al. (2015) the input layer IV is located in the middle compartment and the deep layers V/VI are located in the inner compartment. Analyses are shown both for three anatomically-relevant layer compartments and for four equally-spaced layer compartments.

**Figure 2.**
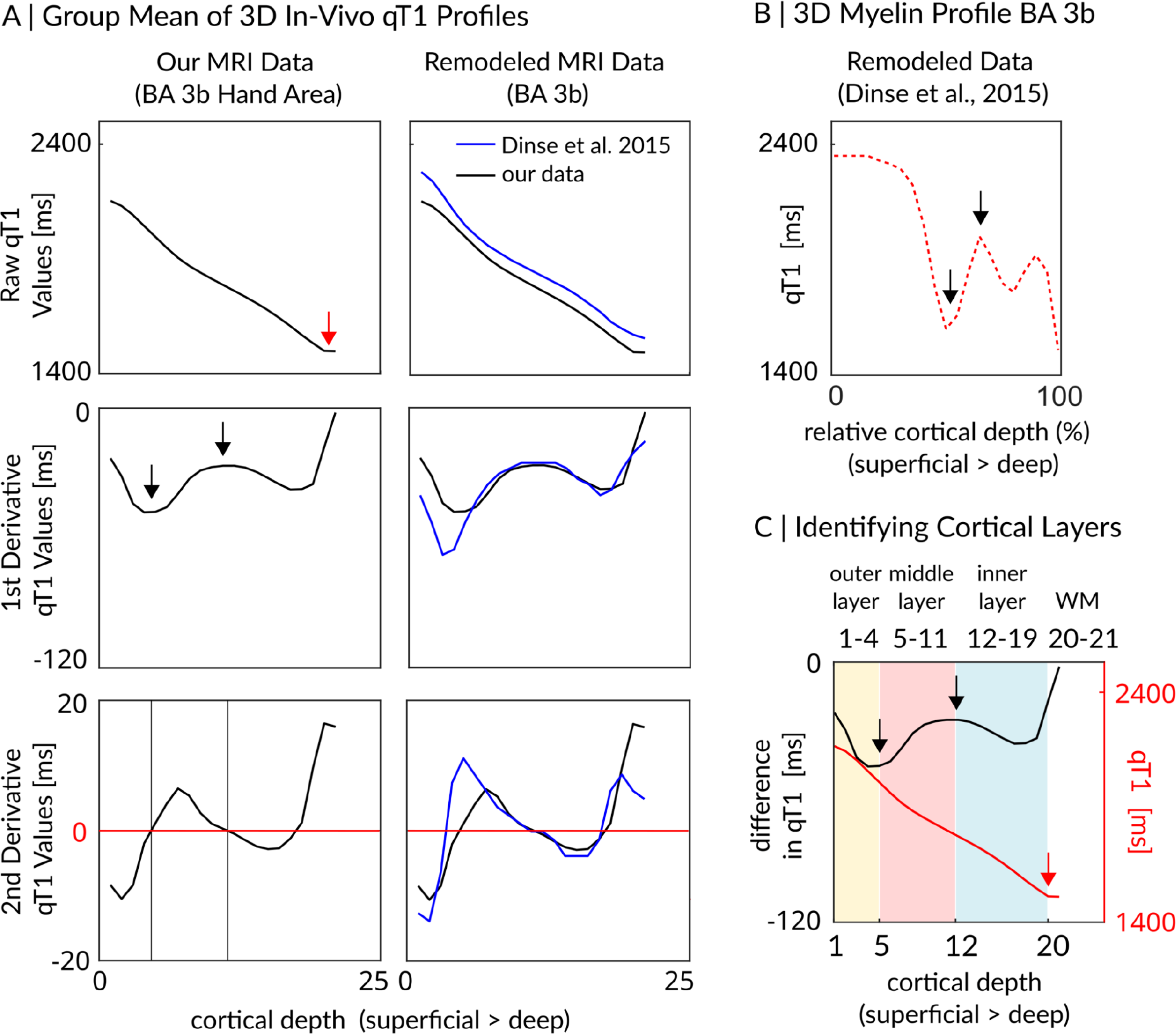
Extraction of anatomically-relevant cortical compartments in BA 3b. (A) Mean qT1 values (y-axis) of the hand representation in BA 3b plotted against 21 cortical depths (x-axis: from superficial to deep). First row: raw qT1 values given in milliseconds (ms) (lower values indicate higher myelin content), second row: first derivative of raw qT1 values, third row: second derivative of raw qT1 values. A red arrow indicates plateauing qT1 values, black arrows indicate the first minimum and the first maximum. The locations of the extrema were estimated based on identifying the zero locations in the second derivative. For comparison, our in-vivo qT1 profiles (black lines) are shown together with remodeled in-vivo qT1 profiles of BA 3b (blue lines, Dinse et al., 2015, second column). (B) Remodeled myelin-related layer profile of BA 3b (red dotted line, Dinse et al., 2015). Black arrows indicate the outer and the inner border of anatomical layer IV according to Dinse et al. (2015). (C) Data-driven approach to identify three anatomically-relevant cortical layers. The middle layer (light pink) was identified based on the first minimum and the first maximum of the first derivative of raw qT1 values (black arrows). Remaining superficial cortical depths were averaged into an outer layer, while remaining deep cortical depths were averaged into an inner layer. Please note that the two deepest cortical depths, where qT1 values stabilized and may reflect white matter (WM, red arrow), were excluded. Please note that we estimated the cortical layers based on *in-vivo* MRI data; they may therefore not correspond exactly to the anatomical layers as identified by histological data.

#### Anatomical Definition of BA 3b

We manually generated subject-specific BA 3b masks of the left hemisphere based on anatomical landmarks in the qT1 images (Geyer, Schleicher, and Zilles, 1999; Yousry et al., 1997), using the software ITK-SNAP (version 3.6.0). Masks were drawn from the edge (beginning of the crown) of the postcentral gyrus to the fundus of the central sulcus (to make sure to include everything of BA 3b), covering the posterior wall of the postcentral gyrus (Geyer, Schleicher, and Zilles, 1999). Only those slices were masked in which the knob-like (axial slices) or the hook-like shape (sagittal slices) of the motor hand area was visible (Yousry et al., 1997). Adjustments were made where required until the surface mask presented homogenous and without holes.

To specifically reduce the effect of possible segmentation errors on surface-based gradient and between-finger septa analyses (explained below), more conservative versions of our manually generated BA 3b masks were created in 15 cases (P01, P02, P03, P04, P07, P08, P10, P11, P13, P14, P15, P16, P19, P20) where structural and/or functional data indicated that we may have sampled a few vertices of BA1. This refinement allowed us to restrict geodesic paths used for gradient and between-finger septa analyses to BA 3b (see **Figure 6** in the results section) and to minimize the influence of neighboring subregions on the results (i.e., lower myelin content in BA 1) (e.g. Glasser et al., 2016; Glasser and Van Essen, 2011; Sanchez-Panchuelo et al., 2014). However, these more conservative masks might also result in missing parts of BA 3b (and were therefore only applied to the gradient and between-finger septa analyses). In line with previous parcellation approaches (Glasser et al., 2016; Glasser and Van Essen, 2011), we identified the border between BA 3b and BA 1 of S1 based on a drop in myelin (increase in qT1 values) along the edge of the rostral bank of the postcentral gyrus. Please note that a clear sharp border cannot be detected in individual myelin-sensitive MR images (Sanchez-Panchuelo et al., 2014), which may be due to the presence of transition zones (Geyer, Schleicher, and Zilles, 1999). Therefore, an exact delineation of S1 subregions cannot not be achieved, which is why an operational definition based on anatomical landmarks extracted from cytoarchitectonic and multimodal parcellation studies was chosen in the present study.

### Functional Data Processing

Distortion correction of opposite polarity EPIs was performed using a point spread function (PSF) mapping (In et al., 2016). To account for differences in the amount of spatial information between the opposite PE EPIs, a weighted combination of the two distortion-corrected images was applied to generate the final, corrected image. The EPI-images of the functional scans were motion corrected to time point = 0. PSF mapping was applied to the motion-corrected images to perform geometrically accurate image reconstruction. Functional time series were slice time corrected to account for differences in image acquisition time between slices using SPM8 (Statistical Parametric Mapping, Wellcome Department of Imaging Neuroscience, University College London, London, UK). Slice-time corrected functional data resulting from phase-encoded protocols were concatenated.

### Registration of Functional Data

Functional time series were semi-manually registered to the MP2RAGE qT1 image using ITK-SNAP (version 3.6.0, non-rigid transformation, 9 degrees of freedom). Resulting registration matrices were applied to the corresponding functional parameter maps (i.e., t-maps, pRF estimates) in a single step (ANTs, version 2.1.0, nearest neighbor interpolation). The inverse of the resulting registration matrices were used to transform individual ROI masks from structural into functional data space (ANTs, version 2.1.0, nearest neighbor interpolation), allowing us to perform ROI analysis on non-smoothed functional data.

### GLM Analysis of Blocked Design Data

We used the general linear model (GLM) as implemented in SPM8 to individually calculate fixed-effects models on the 1st level of the two blocked-design runs (runs 3 and 4 of the experiment, see **Figure 1**). Because each finger was treated individually and independently, BOLD activation driven by each finger’s tactile stimulation was included in the quantification as an independent measure (Kuehn et al., 2018a; Stringer et al., 2014). Each session was modeled with five regressors of interest (stimulation to D1, D2, D3, D4, D5) and allowed the computation of five linear contrast estimates: Touch to D1, D2, D3, D4, and D5 (e.g. the contrast [-1 4 -1 -1 -1] for touch to D2).

### Bayesian pRF Modeling

Population receptive field (pRF) modeling was performed on the phase-encoded fMRI data following the same procedure as introduced by Liu et al. (2021), incorporating the SPM-based BayespRF Toolbox (freely available for download from https://github.com/pzeidman/BayespRF) written for Matlab (SPM12 and Matlab R2017b). We performed a two-stage analysis: First, a 1st level GLM analysis was conducted using SPM12 to prepare the data for pRF modeling by reducing the number of voxel time courses. At this stage, the task regressors were defined. Five regressors were constructed, corresponding to the five fingers of the right hand. Only time-series data that passed a significance threshold of *p*<.05 uncorrected were taken forward for pRF modeling (Zeidman et al., 2018; Puckett et al., 2020). pRF modeling was restricted to BA 3b (using the above described BA 3b masks) to reduce the computing time (note that pRF modeling of one participant takes around 24 hours for the given input data, i.e., it would take several days to compute whole brain data of one participant). pRF modeling was then conducted on a voxel-by-voxel basis to optimize the fit between an estimated waveform and the empirically measured BOLD-response, by modifying the size and position of the pRF model. We thresholded the posterior model probability at > 0.95 (Liu et al., 2021; Puckett et al., 2020; Zeidman et al., 2018). To define the somatosensory space, the dimensions of the 2D matrix were limited to +/- 12.5. Finally, pRF modeling was performed on the inferior-to-superior dimension (x-dimension) of topographic alignment. The minimal pRF size was restricted to 1/10th of the sensory space occupied by a single fingertip, whereas the maximum pRF size was restricted to the equivalence of all five fingers (i.e., 25 units; Liu et al., 2021). A Gaussian pRF profile was chosen as the response function for the pRF analysis (code available for download at https://gitlab.com/pengliu1120/bayesian-prf-modelling.git). The model was characterized by a normal, excitatory distribution with pRF center location (x) and pRF width (i.e., σ, the standard deviation of the Gaussian profile) as parameter estimates. The extracted distance parameter was used to define the digit ROIs (locations of activated voxels for each finger) whereas the extracted width parameter was used as pRF size estimate of activated voxels. Because the stimulus space was one-dimensional, only pRF distance (i.e. center location) and pRF size parameters were analyzed in surface space.

### Localizing Single Fingers in BA 3b

Resulting pRF center location maps were used to locate single finger representations in BA 3b. We applied a “winner-takes-it-all” approach to the pRF center location maps to ensure vertices are sampled only once (i.e. excluding overlapping map areas between finger representations). Vertices of overlapping areas (introduced by splitting pRF maps into five single finger maps and mapping single finger maps onto the inflated surfaces) were exclusively assigned to one area by taking the highest variance (explained by the pRF model) as criterion. The hand area was defined by combining the five fingers to one larger ROI.

### Localizing the Face-Hand Area in BA 3b

A sub-sample (n=16 participants) also underwent functional 7T MRI (gradient-echo EPI sequence of 1.5 mm isotropic resolution with part-brain coverage of M1/S1) while carrying out motor movements of the tongue and the fingers to locate the face-hand area in M1 (Northall et al., 2023). These movement-related data were used to locate the face and hand areas in BA 3b. After the functional data was manually registered to the MP2RAGE qT1 image based on anatomical landmarks (ITK-SNAP version 3.6.0, non-rigid transformation, 9 degrees of freedom), functional activation maps of tongue and finger movements were estimated (peak clusters of t-values, thresholded at p < 0.01 with a minimum cluster size of k=3) using the GLM as implemented in SPM12 (first-level analysis based on contrast estimates for each body part, for details see Northall et al., 2023). A “winner-takes-it-all” approach was applied to the resulting localizers to ensure vertices are sampled only once (i.e., excluding overlapping map areas between body part representations).

### Surface Mapping of Functional Parameters

Registered functional parameters were mapped onto the individual surfaces in anatomical space using the method of closest point (CBS Tools, Bazin et al., 2014, 2001, Surface Mesh Mapping algorithm), and BA 3b masks were applied. To minimize the effect of superficial veins on BOLD signal change, the most superficial 20% of cortical values were disregarded and the mean value of the remaining layers (20–100% cortical depth) were used to perform statistical analyses. Note that no cortical depth-dependent analyses were performed for the functional data.

### Extracting Microstructural Profiles

We sampled qT1, nQSM, pQSM and aQSM values perpendicular to the cortical sheet in 21 different cortical depths from the pRF center locations of the five fingers, and from the face-hand area in BA 3b. Within subjects, extracted values were averaged across vertices (resulting in one value per location/body part and depth). We then calculated the first and the second derivative of resulting qT1 vectors using the gradient function implemented in MATLAB (R2017b). Minima and Maxima of the first derivative were estimated by finding the zero locations in the second derivative.

### Extraction of Structural Gradients

We analyzed qT1, signed QSM (QSM) and aQSM values in relation to geodesic distances to account for local and individual differences in cortical curvature. Geodesic distances and quantitative structural values (qT1, QSM, aQSM) were extracted from BA 3b along the shortest path between the D1 activation peak and the D5 activation peak (19/20 participants, based on blocked design data), using the Dijkstra algorithm as implemented in Pyvista for Python (Sullivan and Kaszynski, 2019). In one case, where D1 and D2 activation maps appeared in reversed inferior-to superior order (P07), the shortest path was calculated between the D2 activation peak and the D5 activation peak. Extracted quantitative values and geodesic distances were used to perform statistical analyses.

### Septa Analysis

To investigate whether between-finger septa also exist in human BA 3b, a surface-based mapping approach combining functional and structural data was used (Kuehn et al., 2017). Layer-specific qT1 values and functional activity (*t*-values generated based on blocked design data) were sampled along predefined paths between neighboring finger representations. In particular, we compared qT1 values extracted from peak locations (maximum t-value of single finger contrast restricted by corresponding pRF map) with qT1 values extracted from border locations (intersection point of t-value vectors sampled along the shortest path between peak locations of neighboring BA 3b finger representations, similar approach to Kuehn et al., 2017). We used the Dijkstra algorithm as implemented in Pyvista for Python (Sullivan and Kaszynski, 2019) to calculate the shortest path between peak locations of neighboring BA 3b finger representations (D1-D2, D2-D3, D3-D4, D4-D5; as well as between D2-D1 and D1-D3 in one case where locations of D1 and D2 finger maps were reversed). qT1 values of n=20 younger adults were therefore sampled from 9 different locations (5 peaks, 4 borders) at 3 different cortical depths (inner, middle, outer).

Additionally, we implemented an analysis that uses multidimensional sampling (inferior-to-superior and anterior-to-posterior) to detect potential low-myelin borders in surface-based qT1 patterns. This analysis is independent of the exact functional crossing vertex and therefore potentially more robust. To realize this approach, we chose the peak activation area of the thumb (as identified using pRF mapping) as a seed region. We then sampled multiple geodesic paths superior and inferior from this seed region, using the Dijkstra algorithm as implemented in Pyvista for Python (Sullivan and Kaszynski, 2019). Start and end points of these paths were extracted along the y-axis of finger activation peaks (D1 and D2), as well as 10 mm below the D1 activation peak (geodesic distance on the shortest path between the D1 activation peak and the face activation peak). This 10 mm distance was estimated based on the expected location of the upper face (in particular the forehead) (Root et al. 2022). Only those vertices that were scattered within one vertex-to-vertex distance (appr. 0.26 mm) around the y-axis were considered. Please note that in two cases (P11, P15), where sampling along the y-axis did not match the underlying surface-based qT1 pattern, start and end points of the geodesic paths were defined along the x-axis. For each participant we sampled five equally-spaced geodesic paths (running from the approximated forehead representation via the thumb to the index finger representation, see **Figure 8A**). Second, we extracted qT1 values from middle cortex depth (where low-myelin borders in BA 3b have been detected previously, Kuehn et al. 2017) and used 3D plotting together with sequential color mapping to visualize the underlying qT1 pattern along the extracted paths (see **Figure 8B**).

### Similarity Analysis

To investigate similarity of finger-specific 3D structural profiles we calculated the Fréchet distance between all possible finger pairs individually for each participant using the Frechet Distance Calculator (version 2.0, Ursell, 2022) as implemented in Matlab (R2022a). The Fréchet distance is a measure of (dis-)similarity between two curves in space which reflects the shortest path length sufficient to traverse both curves from start to finish, while taking the course of the curve into account (Alt and Godau, 1995). When both curves are perfectly aligned the value of the Fréchet distance is zero.

### Statistics

Statistical analyses were computed in R (version 3.4.4, R Core Team, 2022). All sample distributions were analyzed for outliers using boxplot methods and tested for normality using Shapiro-Wilk’s test. In case of non-normal data, non-parametric or robust tests were conducted. For all tests, the significance level was set to p = 0.05. Bonferroni-corrected significance levels were applied for multiple testing to correct for family-wise error accumulation. Pearson’s correlation coefficient r was calculated as an effect size estimator for Student t-tests (Field et al. 2012). We applied Cohen’s criteria of 0.3 and 0.5 for a medium and large effect, respectively. Eta-squared (*η²*) was calculated as an effect size estimator for ANOVAs. We applied two-sided tests, except for clear one-sided hypotheses (i.e. testing for low-myelin borders).

To test for the existence of inferior-to-superior structural gradients in the human BA 3b hand area (potential linear relationships between geodesic distances and quantitative structural values), we used multilevel linear regression models to predict layer-specific qT1, QSM and aQSM values from normalized geodesic distances (fixed effect) while controlling for the effect of individuals (random effect with random intercept and random slope). To conduct this analysis, we used the lme function (method maximum likelihood) from the nlme package (version 3.1-155, Pinheiro et al., 2022) as implemented in R (version 3.4.4, R Core Team, 2022).

In addition, we fitted Bayesian linear multilevel models using the brm function of the brms package (version 2.18.0, Bürkner 2017, Bürkner 2018, Carpenter et al. 2017) as implemented in R (version 4.2.2, R Core Team, 2022). We used the same fixed and random effects as described before for the frequentist approach. The prior distribution of the intercept for layer-dependent qT1 was informed by previous 7T MRI studies (Dinse et al. 2015, Kuehn et al. 2017). Based on this information, we chose normal prior distributions with different means and standard deviations for different layers (outer: mean = 2058, standard deviation = 483, middle: mean = 1770, standard deviation = 294, inner: mean = 1703, standard deviation = 183). Previously reported values were averaged and rounded to whole numbers, standard deviations were multiplied by 3 to allow variation while accounting for possible outliers in the data. The prior for the effect of geodesic distance on qT1 was set to normal (mean = 0 for all layers, same standard deviations as reported for the intercepts). QSM and aQSM priors were also informed by previous studies (Acosta-Cabronera et al. 2018, Donatelli et al. 2022). Signed QSM (QSM) values were allowed to vary between -0.125 and 0.125, aQSM values were allowed to vary between 0 and 0.125. The mean of the prior distribution was set to 0.5 times the possible value range, the standard deviation to 0.25 times the possible value range. We chose normal prior distributions for the intercept and the effect of geodesic distance on QSM (mean = 0, standard deviation = 0.06) and aQSM (mean = 0.06, standard deviation = 0.03) for all layers. Because we were interested in a linear relationship between structural measures (qT1, QSM, aQSM) and geodesic distance, the Gaussian family with the identity link was chosen as response distribution (Bürkner 2017). All models were fitted using 4 chains with 2000 iterations each, of which the first 1000 were used to calibrate the sampler, resulting in a total of 4000 posterior samples (Bürkner 2017).

To compare skewness and kurtosis of microstructure profiles between fingers (D1, D2, D3, D4, D5) as well as to compare Fréchet distances of microstructure profiles between pairs of neighboring fingers (D1-D2, D2-D3, D3-D4, D4-D5), one-way repeated-measures ANOVAs on qT1, nQSM, pQSM and aQSM were computed given normality. In case of non-normal data, we conducted robust repeated-measures ANOVAs based on 20%-trimmed means which were performed using the rmanova function from the WRS2 package (version 1.1-3, Mair and Wilcox, 2020) in R.

To test for layer-specific differences in qT1, nQSM, pQSM and aQSM between finger representations, we calculated two-way repeated-measures ANOVAs with layer (inner, middle, outer) and location (D1, D2, D3, D4, D5) as within-subjects factors. In case of sphericity violations, Greenhouse-Geisser-corrected results (when violations to sphericity were large, i.e. ε < .75) or Huynd-Feldt-corrected results (when violations to sphericity were small, i.e. ε >= .75) were computed. Post-hoc tests were performed as two-tailed paired-samples tests. Non-parametric Wilcoxon signed rank tests were additionally calculated where conditions of the parametric one-sample t-test were violated. For sample distributions containing extreme outliers (values above the third quartile plus three times the interquartile range or values below the first quartile minus three times the interquartile range), two-way repeated-measures ANOVAs were computed both with and without extreme outliers included. Please note that in order to offer data that can more easily be compared to previous studies (Kuehn et al. 201, Tardiff et al., 2015), we also conducted the analyses using the equally-spaced layer definition that, however, may not reflect anatomically-relevant cortical compartments.

To test for layer-specific differences in qT1, nQSM, pQSM and aQSM between the face and the hand representation, we calculated two-way repeated-measures ANOVAs with layer (inner, middle, outer) and body part (face, hand) as within-subjects factors using the same correction methods and post-hoc tests as described above.

To investigate whether between-finger septa also exist in human BA 3b, we performed layer-wise comparisons on peak-to-border differences of qT1 values by applying Bayesian paired-samples t-tests (resulting in 12 tests given four different peak-to-border conditions (D1-D2, D2-D3, D3-D4, D4-D5) and three different cortical depths). For this purpose, qT1 values were averaged across neighboring finger representations for each peak-to-border condition (i.e., D1-D2 average, D2-D3 average, D3-D4 average, D4-D5 average). The JASP software package (version 0.8.1.2, JASP Team, 2017) was used to calculate Bayesian paired-samples t-tests with location (peak, border) as within-subjects factor. We used a Bayes factor favoring the alternative hypothesis. The default Cauchy-scaled prior of 0.707 was selected instead of modeling the expected effect size. However, we simulated the effect of the prior on the Bayes factor for a wider range of prior width to estimate how robust the conclusions were to the chosen prior.

## Results

### Cortical 3D Profiles and Layer Definition

To study the 3D structural architecture of the human hand representation in BA 3b in reference to functional topography, we used quantitative T1 (qT1) as a proxy for cortical myelin (Stüber et al., 2014, Dinse et al., 2015) and quantitative susceptibility maps (QSM) as proxies for iron (positive values, pQSM), diamagnetic tissue substances (negative values, nQSM) (Langkammer et al., 2012, Zheng et al., 2013, Hametner et al., 2018), and overall mineralization (absolute values, aQSM) (Betts et al., 2016). In addition to anatomical scans, also functional scans were acquired when participants were stimulated at the fingertips of their right hand by an automated piezoelectric stimulator, using both phase-encoded and blocked designs. We focussed on investigating left BA 3b (contralateral to where tactile stimulation was applied). pRF center locations as revealed by Bayesian pRF modeling were used to locate each finger in BA 3b (thumb: D1, index finger: D2, middle finger: D3, ring finger: D4, little finger: D5) (Liu et al., 2021).

When extracting qT1, nQSM, pQSM and aQSM values from the hand area in BA 3b, we obtained precise intracortical contrasts (see **Figure 3A**). As expected (Stüber et al., 2014; Dinse et al., 2015), qT1 values decrease from superficial to deep cortical depths, reflecting high myelin close to the white matter (WM). This intracortical gradient from low to high myelin is interrupted in the upper half of the profile, an area known to contain high myelin content (i.e., Baillarger band in anatomical layer IV) (Vogt and Vogt, 1919; Dinse et al., 2015). Based on previous *in vivo ex vivo* validation work (Dinse et al., 2015), we used the local rate of change of qT1 values to define three anatomically-relevant cortical compartments (here referred to as ‘outer layer’, ‘middle layer’ and ‘inner layer’, see **Figure 3B**, **Figure 3C**), where input layer IV is assumed to be located in the middle layer, and layers V and VI are assumed to be located in the inner layer. Note that the upper peak of the pQSM profile (which showed a top-hat distribution shaped by a plateau with a double peak) is located in what we define as middle layer (containing layer IV), whereas the lower peak is located in what we define as inner layer (containing layer V). With respect to nQSM, we observed a u-shaped profile with values closest to zero in the very middle of cortical depths, which is interrupted by a small plateau in the upper half of the profile, where we expected layer IV to be located. Multi-contrast mapping therefore confirmed our qT1-based layer definition. Statistical results are shown both for three layers (outer layer, middle layer, inner layer) and for four equally-spaced layers to facilitate comparing our data to previous publications (Kuehn et al., 2017, Tardiff et al., 2015).

**Figure 3.**
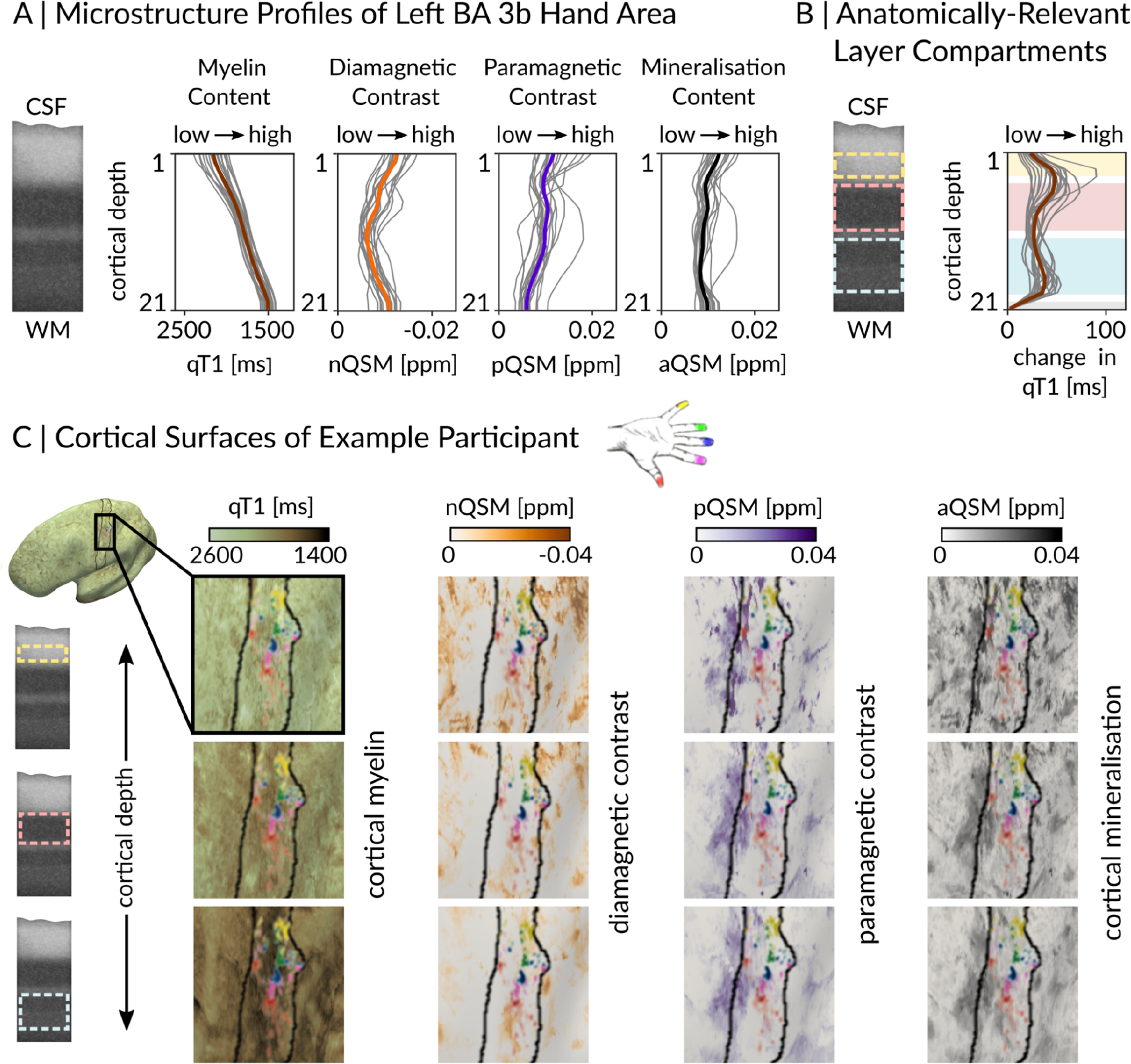
Cortical 3D Profiles and Layer Definition. (A) Microstructure profiles of the left (contralateral to where stimulation was applied) hand representation in BA 3b show distinct cortical depth-dependent profiles. Quantitative T1 (qT1, reflecting myelin, n=20), negative QSM (nQSM, reflecting diamagnetic tissue contrast (e.g., calcium), n=18), positive QSM (pQSM, reflecting paramagnetic tissue contrast (iron), n=18) and absolute QSM (aQSM, overall mineralization, n=18) were sampled at 21 different cortical depths (Kuehn et al., 2017) where depth 1 is located closest to cerebrospinal fluid (CSF) and depth 21 is located closest to white matter (WM). Myelin staining was remodeled according to Dinse et al. (2015). (B) Anatomically-relevant cortical compartments (cream: outer layer, light pink: middle layer, light blue: inner layer) were defined based on minima and maxima of the rate of change in qT1 (first derivative of cortical depth-dependent qT1 calculated as central difference between two neighboring sampling points) that are assumed to reflect the heavily myelinated Bands of Baillarger. (C) qT1, nQSM, pQSM and aQSM values (from left to right) of the left hand area in BA 3b in each of the layers (top row: outer layer, middle row: middle layer, bottom row: inner layer) for one example participant. The population receptive field (pRF) center is plotted on top to show individual finger representations from which data were extracted (red: thumb, magenta: index finger, blue: middle finger, green: ring finger, yellow: little finger).

#### Absence of Inferior-to-Superior Structural Gradients in the BA 3b Hand Area

First, we investigated whether there is a systematic structural gradient within the hand area of BA 3b (e.g. higher myelin in superior compared to inferior parts of the hand area). To this end, we extracted layer-specific qT1, signed QSM (QSM), and aQSM values along geodesic paths from inferior to superior and calculated multilevel linear regression models (fixed effect: normalized geodesic distance, random effect: individual). None of the tested models were significant (see **Table 1** and **Figure 4**), indicating an absence of a systematic inferior-to-superior structural gradient in the hand area in BA 3b, which was also confirmed for four equally-spaced layers (see **Table 1**), and by calculating Bayesian multilevel models (see **Table 2**).

**Figure 4.**
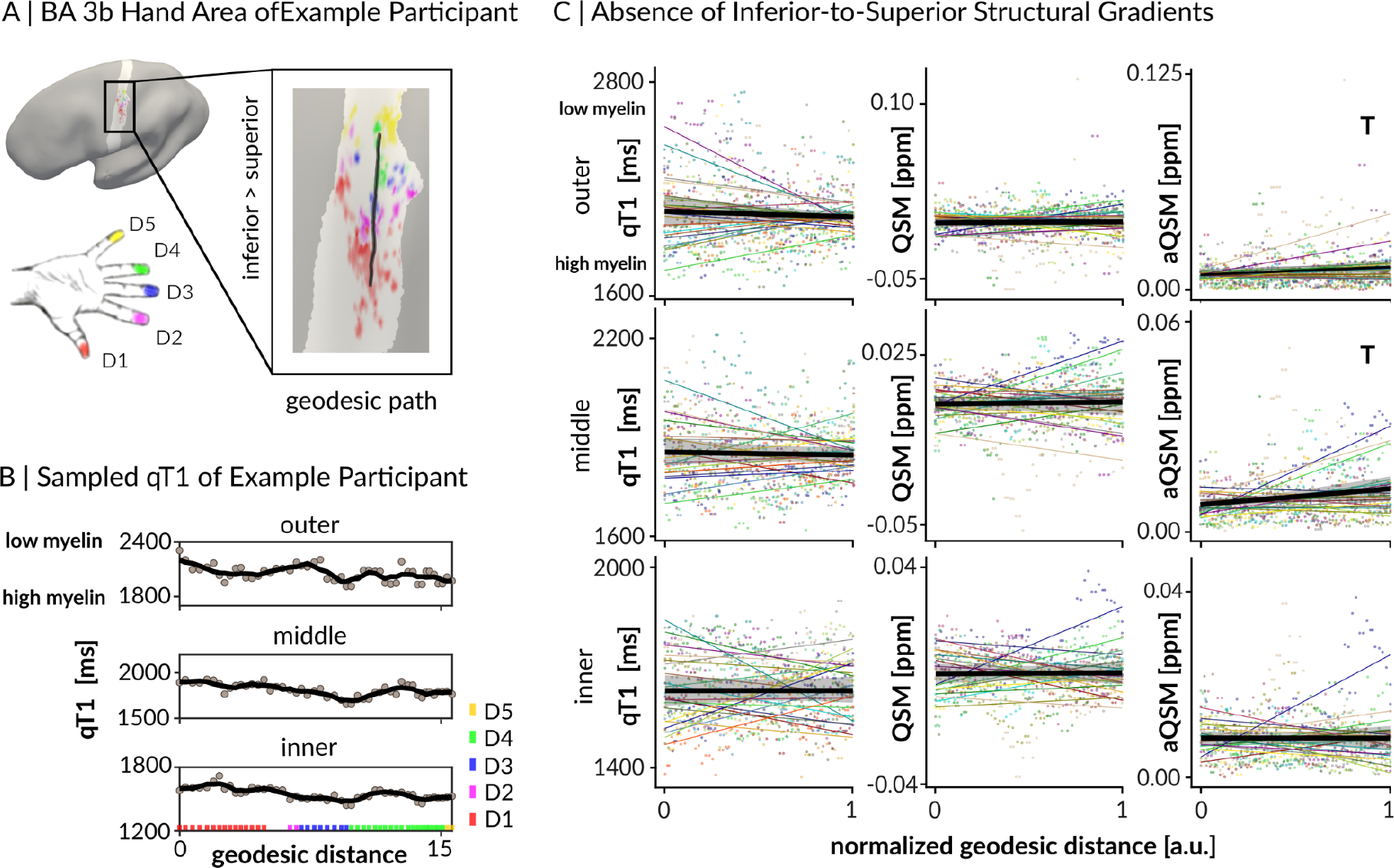
Absence of inferior-to-superior structural gradient in BA 3b hand area. (A) Using surface-based analyses, structural gradients of quantitative T1 (qT1), signed QSM (QSM) and absolute QSM (aQSM) were extracted from the full fingertip map (= hand area) of left BA 3b (light gray area). Values were extracted from inferior to superior along the geodesic path (black line). Different colors on the schematic drawing of the hand indicate different finger representations on the cortical surface. (B) qT1 values of one example participant sampled from inferior to superior vertices along the geodesic path for anatomically-relevant layers. Black lines represent local k-point mean values calculated over a sliding window of length k (15% of all data points) across neighboring vertices, light gray values represent individual data points, and different colors indicate different fingers. (C) Layer-dependent mean gradients (black lines) of qT1 values (in ms), QSM values (in ppm) and aQSM values (in ppm) are shown together with 95% confidence intervals (gray ribbons), individual data points (colored dots) and individual trend lines (colored lines). There were no significant results. Trends above Bonferroni-corrected threshold of p=0.0167 (correcting for three tests per structural variable) are marked by a T.

**Table 1.** Statistical results of frequentist gradient analysis using multilevel linear regression. None of the tested layer-specific structural measures (qT1 = quantitative T1, QSM = signed QSM, aQSM = absolute QSM) showed a significant linear relationship with geodesic distances. Structural gradients were tested for anatomically-relevant layers (outer, middle, inner) and for equally-spaced layers (sf = superficial, om = outer-middle, im = inner-middle, dp = deep). Bonferroni-corrected significance levels of p = 0.0167 for anatomically-relevant layers (correcting for 3 tests per structural variable) and p = 0.0125 for equally-spaced layers (correcting for 4 tests per structural variable) were applied. Trends above Bonferroni-corrected significance levels are marked by a T. Results are reported as slope (β), Student’s t (t), p-value (p) and 95% confidence intervals (95% CI). Geodesic distances were normalized.

| anatomically-relevant layers |  |  |  |  | equally-spaced layers |  |  |  |  |
| --- | --- | --- | --- | --- | --- | --- | --- | --- | --- |
| model | qT1 ~ geodesic distance, n=20 |  |  |  | model | qT1 ~ geodesic distance, n=20 |  |  |  |
| layer | $\beta$ | $t$ | $p$ | 95% CI | layer | $\beta$ | $t$ | $p$ | 95% CI |
| outer | -32.70 | -0.68 | 0.50 | -126.88, 61.47 | sf | -22.39 | -0.76 | 0.45 | -80.28, 35.50 |
| middle | -8.87 | -0.40 | 0.69 | -51.88, 34.15 | om | -6.13 | -0.27 | 0.78 | -49.86, 37.60 |
| inner | 3.01 | 0.10 | 0.92 | -57.04, 63.07 | im | 2.24 | 0.09 | 0.93 | -47.40, 51.88 |
|  |  |  |  |  | dp | 2.54 | 0.07 | 0.94 | -64.17, 69.25 |
| model | QSM ~ geodesic distance, n=18 |  |  |  | model | QSM ~ geodesic distance, n=18 |  |  |  |
| layer | $\beta$ | $t$ | $p$ | 95% CI | layer | $\beta$ | $t$ | $p$ | 95% CI |

**Table 1. Statistical results of frequentist gradient analysis using multilevel linear regression.**
| outer | 0.001 | 0.27 | 0.79 | -5.9<sup>-3</sup>, 7.8<sup>-3</sup> | sf | 4.6<sup>-4</sup> | 0.13 | 0.89 | -6.3<sup>-3</sup>, 7.3<sup>-3</sup> |
| --- | --- | --- | --- | --- | --- | --- | --- | --- | --- |
| middle | 0.001 | 0.29 | 0.77 | -5.6<sup>-3</sup>, 7.5<sup>-3</sup> | om | 0.001 | 0.42 | 0.68 | -5.3<sup>-3</sup>, 8.2<sup>-3</sup> |
| inner | 4.1<sup>-4</sup> | 0.13 | 0.89 | -5.6<sup>-3</sup>, 6.4<sup>-3</sup> | im | 0.001 | 0.40 | 0.69 | -5.0<sup>-3</sup>, 7.5<sup>-3</sup> |
|  |  |  |  |  | dp | -4.6<sup>-4</sup> | -0.14 | 0.89 | -6.9<sup>-3</sup>, 6.0<sup>-3</sup> |
| model aQSM ~ geodesic distance, n=18 |  |  |  |  | model aQSM ~ geodesic distance, n=18 |  |  |  |  |
| layer | $\beta$ | $t$ | $p$ | 95% CI | layer | $\beta$ | $t$ | $p$ | 95% CI |
| outer | 0.005 | 1.94 | 0.05 T | -4.3<sup>-5</sup>, 9.3<sup>-3</sup> | sf | 0.006 | 3.03 | 0.03 T | 2.2<sup>-3</sup>, 1.0<sup>-2</sup> |
| middle | 0.005 | 2.12 | 0.04 T | 3.2<sup>-4</sup>, 8.9<sup>-3</sup> | om | 0.004 | 1.53 | 0.13 | -1.0<sup>-3</sup>, 8.3<sup>-3</sup> |
| inner | -2.8<sup>-5</sup> | -0.02 | 0.99 | -3.6<sup>-3</sup>, 3.5<sup>-3</sup> | im | 7.9<sup>-5</sup> | 0.03 | 0.97 | -4.7<sup>-3</sup>, 4.8<sup>-3</sup> |
|  |  |  |  |  | dp | -7.8<sup>-5</sup> | -0.04 | 0.96 | -3.5<sup>-3</sup>, 3.3<sup>-3</sup> |

**Table 2.** Statistical results of Bayesian gradient analysis. None of the tested layer-specific structural measures (qT1 = quantitative T1, QSM = signed QSM, aQSM = absolute QSM) showed a linear relationship with geodesic distances (i.e., BF_10_ < 1). Structural gradients were tested for anatomically-relevant layers (outer, middle, inner). Results are reported as estimated means of the population-level effect of geodesic distance on qT1 (β), 95% credible interval based on quantiles (95% CI), and Bayes factor favoring the alternative hypothesis (BF_10_). A Bayes factor of 1 or greater represents evidence in favor of the alternative hypothesis (H1: linear gradient is present), and a Bayes factor < 1 represents evidence in favor of the null hypothesis (H0: linear gradient is absent). The convergence diagnostic for Hamiltonian Monte Carlo chains indicated convergence for all models (Rhat ≤ 1.01).

| model | qT1 ~ geodesic distance<br>n=20 |  |  | QSM ~ geodesic distance<br>n=18 |  |  | aQSM ~ geodesic distance<br>n=18 |  |  |
| --- | --- | --- | --- | --- | --- | --- | --- | --- | --- |
| layer | $\beta$ | 95% CI | $BF_{10}$ | $\beta$ | 95% CI | $BF_{10}$ | $\beta$ | 95% CI | $BF_{10}$ |
| outer | -31.7 | -128.3,<br>61.7 | 0.12 | 8.0<sup>-4</sup> | -7.4<sup>-3</sup>, 8.5<sup>-3</sup> | 0.063 | 4.8<sup>-3</sup> | -4.0<sup>-4</sup>, 1.0<sup>-2</sup> | 0.45 |
| middle | -9.5 | -54.9, 34.7 | 0.08 | 7.0<sup>-4</sup> | -7.3<sup>-3</sup>, 8.1<sup>-3</sup> | 0.056 | 4.6<sup>-3</sup> | -1.0<sup>-4</sup>, 9.2<sup>-3</sup> | 0.44 |
| inner | 1.2 | -58.1, 62.5 | 0.16 | 5.0<sup>-4</sup> | -6.3<sup>-3</sup>, 7.3<sup>-3</sup> | 0.056 | 1.3<sup>-5</sup> | -4.1<sup>-3</sup>, 4.2<sup>-3</sup> | 0.07 |

#### Absence of Low-Myelin Borders between Finger Representations in BA 3b

Next, we targeted the question on the presence or absence of low-myelin borders between finger representations in human BA 3b by comparing qT1 values sampled from the functional peak (highest t-value) of each finger representation with qT1 values sampled from the functional border between neighboring finger representations. The shortest path was estimated using the surface-based Dijkstra-algorithm (see **Figure 5A**), which is analogous to the way low-myelin borders between the hand and the face were detected (Kuehn et al., 2017). We calculated layer-specific Bayesian probabilities (Bayes factors) as a relative measure of similarity for all neighboring finger pairs (D1-D2, D2-D3, D3-D4, D4-D5). Our results revealed that the alternative hypothesis (finger peak qT1 values < border qT1 values) was predicted a maximum 0.86 times better than the null hypothesis (no difference in qT1 values between peak and border) in any layer (see **Table 3**, **Figure 5E**). This provides evidence in favor of the null hypothesis of no difference, indicating an absence of low-myelin borders between finger representations in human BA 3b.

**Figure 5.**
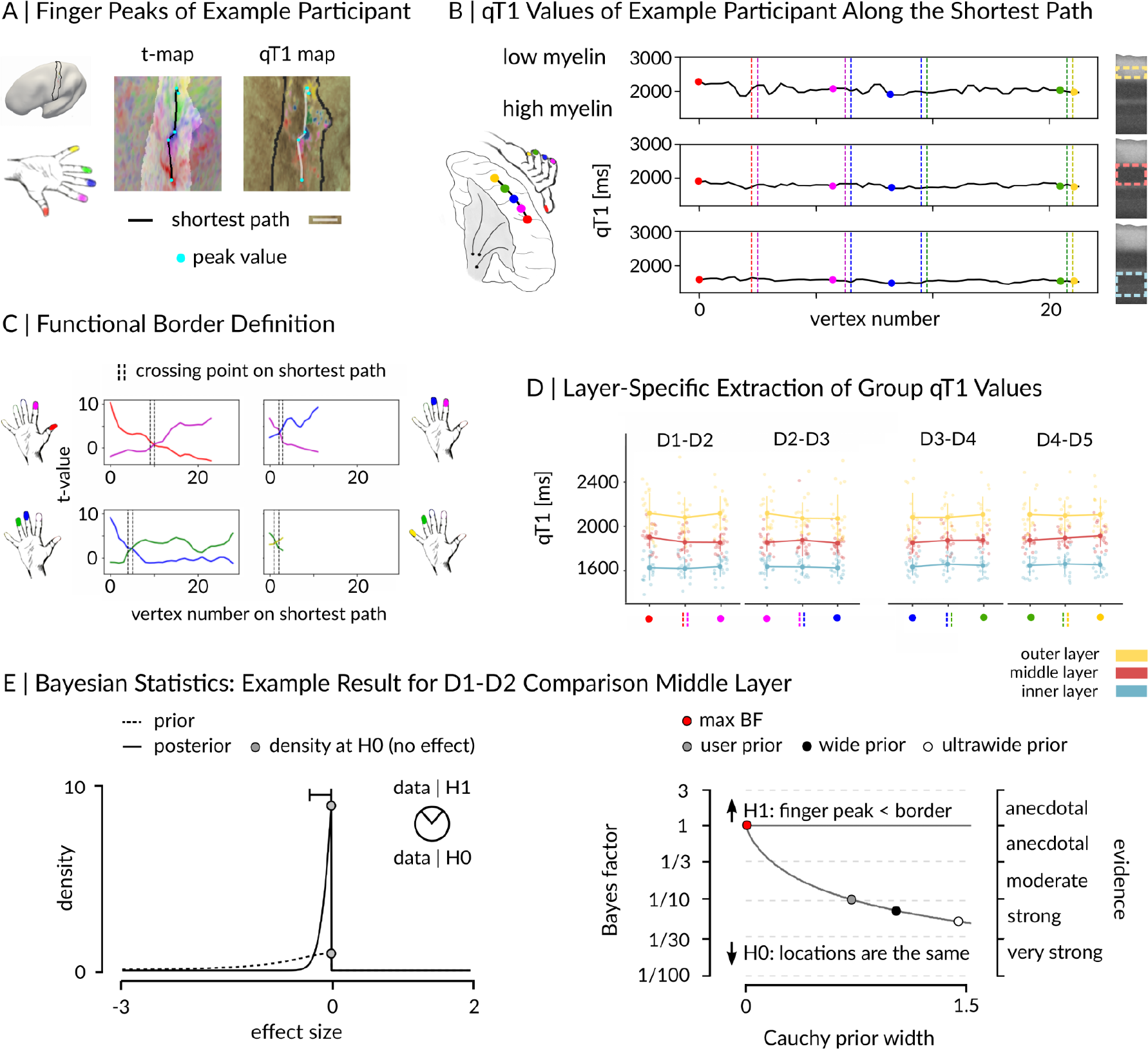
Absence of low-myelin borders between finger representations in human BA 3b. (A) Locations of finger peak values (cyan dots) and estimated shortest paths between finger peaks (based on Dijkstra algorithm) within left BA 3b (light area in t-map, solid black line in qT1 map) are shown together with mapped t-values (five contrasts for five fingers, e.g. D1 (red) and qT1 values (darker areas reflect lower qT1 values, i.e. higher myelin content). (B) qT1 values were sampled vertex-wise along the shortest paths between finger peak locations (colored dots) in each layer (yellow: outer, red: middle, light blue: inner). Dotted lines indicate borders between neighboring fingers. (C) Locations of functional borders between fingers were defined as the vertex on the shortest paths where the t-value functions of neighboring finger representations were equal (= crossing point, dashed black lines). (D) qT1 values were sampled from finger peak and border locations (see B) in each layer resulting in 12 comparisons. Colored dots represent individual data, group means are plotted in bold. Whiskers represent the SEM. (E) Example of Bayesian Statistics. A Bayesian paired-sample t-test was calculated to test the peak-to-boder difference in qT1 between D2 and D3 in the middle layer. Prior distributions are shown as dashed lines, posterior distributions as solid lines. Gray dots indicate the height of the curves (density) at the null hypothesis (H0). A Bayes factor of 1 or greater represents evidence in favor of the alternative hypothesis (H1: finger peak < border), and a Bayes factor below 1 represents evidence in favor of the H0 (no difference between finger peak and border). The higher dot on the posterior distribution indicates that the Bayes factor supports the H0 (locations, i.e. finger peak and border, are the same).

**Table 3.**
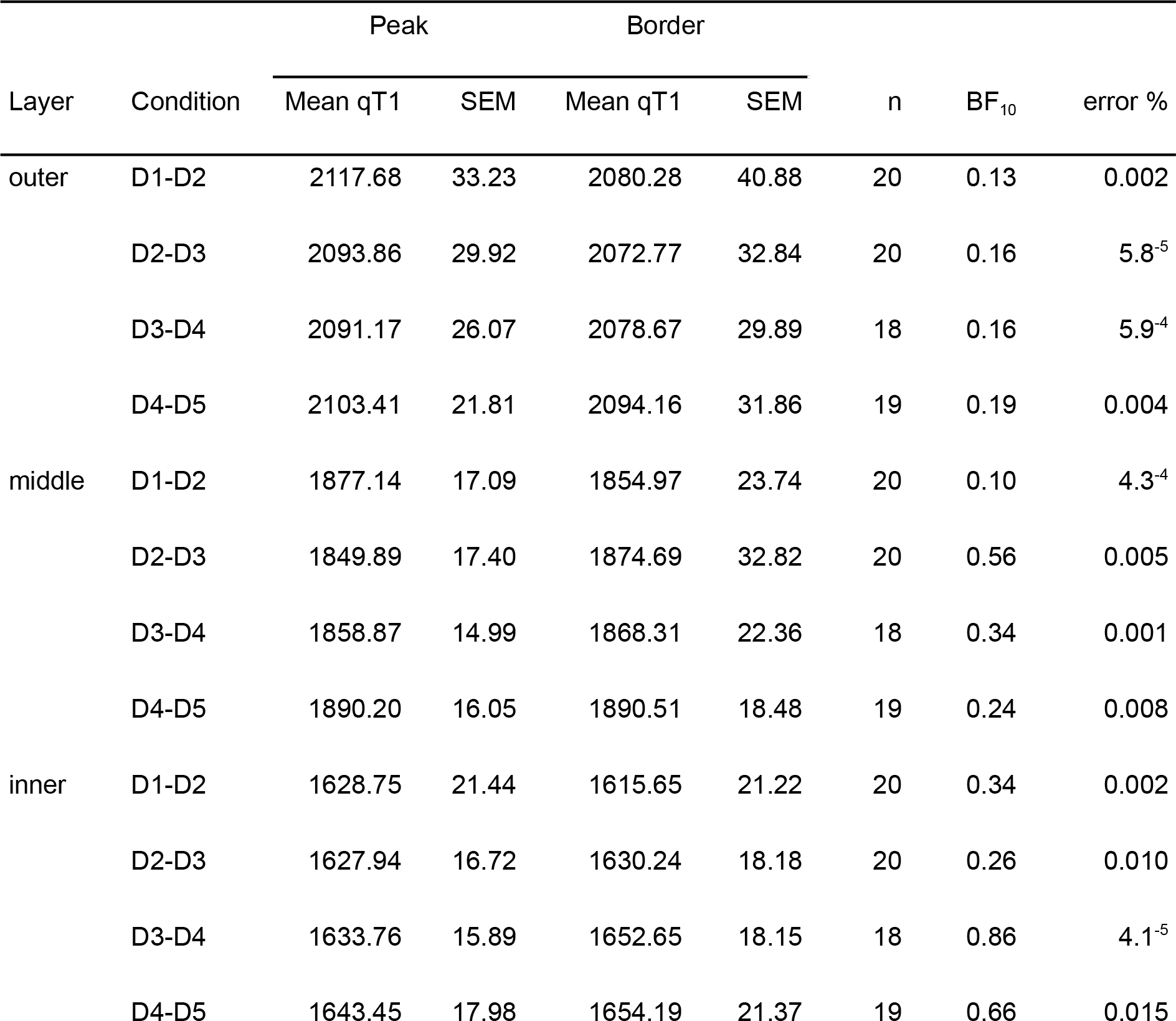
Descriptives and Bayesian statistics of between-finger low-myelin border analysis. Shown are mean qT1 values given in milliseconds and standard errors of the mean (SEM), the number of observations (n), Bayes factors (BF) and errors associated with the Bayes factor given in % (error %). The Bayes factor represents a relative measure of similarity of qT1 values for all finger pairs (D1-D2, D2-D3, D3-D4, D4-D5) at each layer (outer, middle, inner). A Bayes factor of 1 or greater represents evidence in favor of the alternative hypothesis (finger peak qT1 < border qT1), and a Bayes factor below 1 represents evidence in favor of the null hypothesis (no difference between finger peak qT1 values and border qT1 values). One-sided Bayesian t-tests were calculated to test the alternative hypothesis.

| Layer | Condition | Peak |  | Border |  | n | BF<sub>10</sub> | error % |
| --- | --- | --- | --- | --- | --- | --- | --- | --- |
|  |  | Mean qT1 | SEM | Mean qT1 | SEM |  |  |  |
| outer | D1-D2 | 2117.68 | 33.23 | 2080.28 | 40.88 | 20 | 0.13 | 0.002 |
|  | D2-D3 | 2093.86 | 29.92 | 2072.77 | 32.84 | 20 | 0.16 | 5.8<sup>-5</sup> |
|  | D3-D4 | 2091.17 | 26.07 | 2078.67 | 29.89 | 18 | 0.16 | 5.9<sup>-4</sup> |
|  | D4-D5 | 2103.41 | 21.81 | 2094.16 | 31.86 | 19 | 0.19 | 0.004 |
| middle | D1-D2 | 1877.14 | 17.09 | 1854.97 | 23.74 | 20 | 0.10 | 4.3<sup>-4</sup> |
|  | D2-D3 | 1849.89 | 17.40 | 1874.69 | 32.82 | 20 | 0.56 | 0.005 |
|  | D3-D4 | 1858.87 | 14.99 | 1868.31 | 22.36 | 18 | 0.34 | 0.001 |
|  | D4-D5 | 1890.20 | 16.05 | 1890.51 | 18.48 | 19 | 0.24 | 0.008 |
| inner | D1-D2 | 1628.75 | 21.44 | 1615.65 | 21.22 | 20 | 0.34 | 0.002 |
|  | D2-D3 | 1627.94 | 16.72 | 1630.24 | 18.18 | 20 | 0.26 | 0.010 |
|  | D3-D4 | 1633.76 | 15.89 | 1652.65 | 18.15 | 18 | 0.86 | 4.1<sup>-5</sup> |
|  | D4-D5 | 1643.45 | 17.98 | 1654.19 | 21.37 | 19 | 0.66 | 0.015 |

To align our approach as closely as possible with detection approaches of low-myelin borders in monkey species (e.g. Qi and Kaas, 2004), we explored individual qT1 maps of the middle layer visually (i.e., where low-myelin borders should be located, Kuehn et al., 2017). Stripe-like architectures within the hand area (similar to those described in BA 3b of macaque monkeys, Qi and Kaas, 2004) could be observed in 10/20 participants in the following locations (see **Figure 6**): in 2/20 participants between D1 and D2 (P01, P08), in 7/20 participants between D2 and D3 (P01, P10, P11, P15, P16, P18, P20), in 1/20 participants between D2 and D4 (P10), in 2/20 participants between D3 and D4 (P15, P17), and in 2/20 participants between D3 and D5 (P12, P16). Therefore, no systematic stripe-like representation (of potential low-myelin borders separating individual fingers) could be identified between specific finger pairs. But, as expected, low-myelin borders were clearly visible between the hand and the face areas in all participants (except P04, see **Figure 7**).

**Figure 6.**
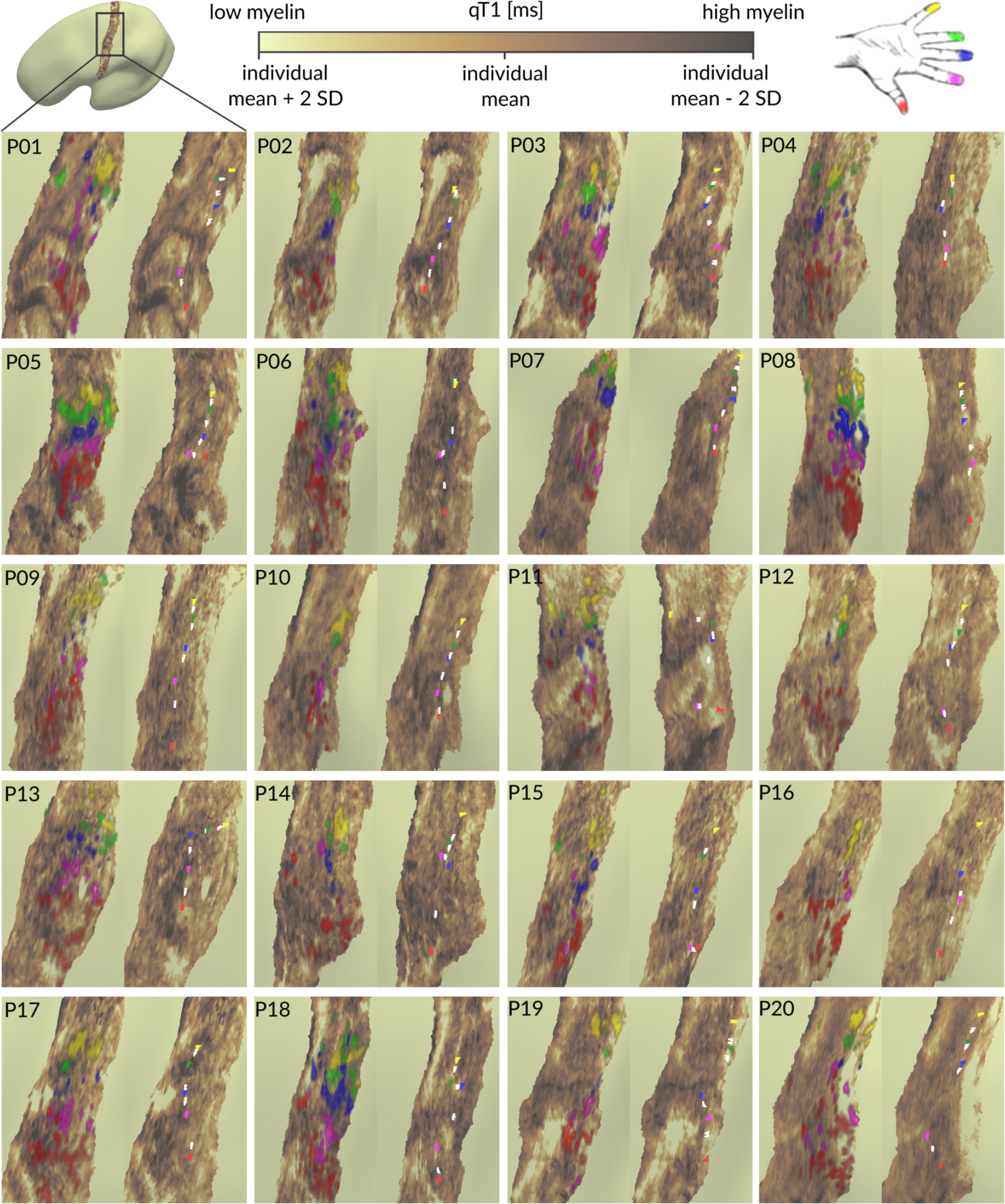
Individual qT1 maps of human BA 3b fingertip representations. Cortical qT1 values extracted from the middle layer of left BA 3b (contralateral to where stimulation was applied) are shown together with pRF center location maps (differently colored areas indicating different functional finger representations), finger activation peaks (colored dots indicating highest t-values, red: thumb, magenta: index finger, blue: middle finger, green: ring finger, yellow: little finger) and functional borders between fingers (white dots indicating crossing point of t-value function of neighboring finger representations). qT1 values are given in milliseconds (ms), value ranges are thresholded within individual means +/- 2 standard deviations (SD). Areas of high myelin content (low qT1 values) are colored in dark brown, while areas of low myelin content (high qT1 values) are colored in olive green.

**Figure 7.**
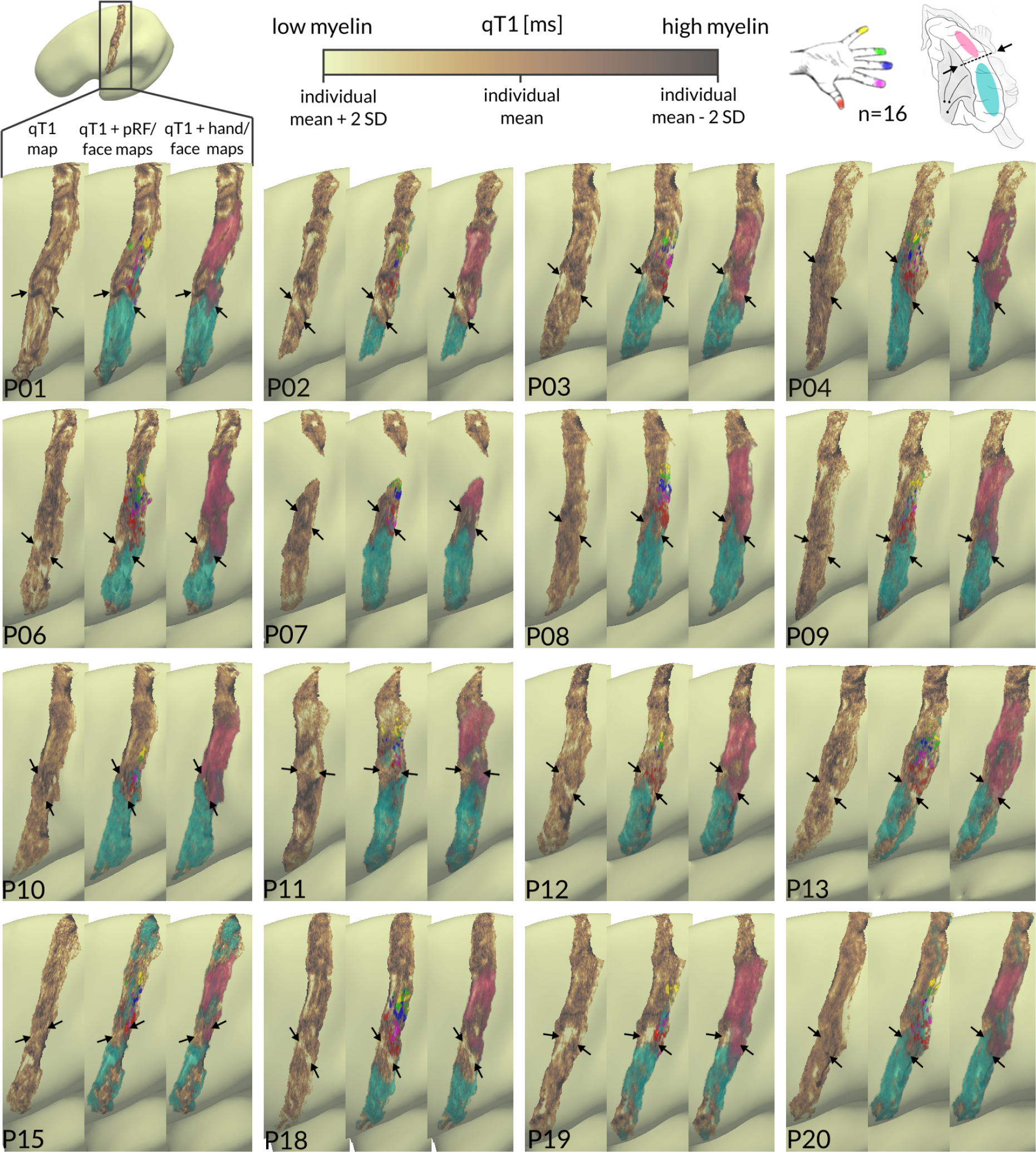
Individual qT1 maps of BA 3b show low-myelin borders between the face and hand areas. Cortical qT1 values extracted from the middle layer of left BA 3b (contralateral to where stimulation was applied or to where movement was carried out) are shown together with pRF center location maps of individual fingers (red: thumb, magenta: index finger, blue: middle finger, green: ring finger, yellow: little finger), functional face activation maps (clusters of highest t-values (p<0.01) colored in cyan) and functional hand activation maps (clusters of highest t-values (p<0.01) colored in light pink). Black arrows indicate low-myelin borders between face and hand representations in BA 3b. qT1 values are given in milliseconds (ms), value ranges are thresholded individually within the mean +/- 2 standard deviations (SD). Areas of high myelin content (lower qT1 values) are colored in dark brown.

Moreover, we carried out an additional analysis that uses multidimensional sampling (inferior-to-superior and anterior-to-posterior) to detect potential low-myelin borders in surface-based qT1 patterns that is independent of the exact functional crossing vertex. qT1 values were sampled from middle cortical depth (where low-myelin borders in BA 3b have been previously detected, Kuehn et al., 2017) along multiple extracted geodesic paths (running from a seed region (D1) inferior to the upper face region and superior to the D2 region in BA 3b, see **Figure 8A**). Using 3D data extraction together with sequential color mapping revealed, as expected, systematic low-myelin patterns (represented by lighter colors in **Figures 8B, 8C**) between the upper face and the thumb region in 14/16 participants (P02, P03, P06, P07, P08, P09, P10, P12, P13, P15, P18, P20, and presumably P11 and P19 see below). These patterns are not only indicative of the location of the hand-face border but also of its shape (when comparing patterns of light-colored qT1 segments with the appearance of low-myelin regions on individual cortical surfaces, shown next to the 3D plots in **Figures 8B, 8C**). For example, in some cases, the light-colored segments of the different paths (indicative of low-myelin regions) form a u-shape (P02, P03), whereas in other cases they appear in a hook-like shape (P08, P10).

**Figure 8.**
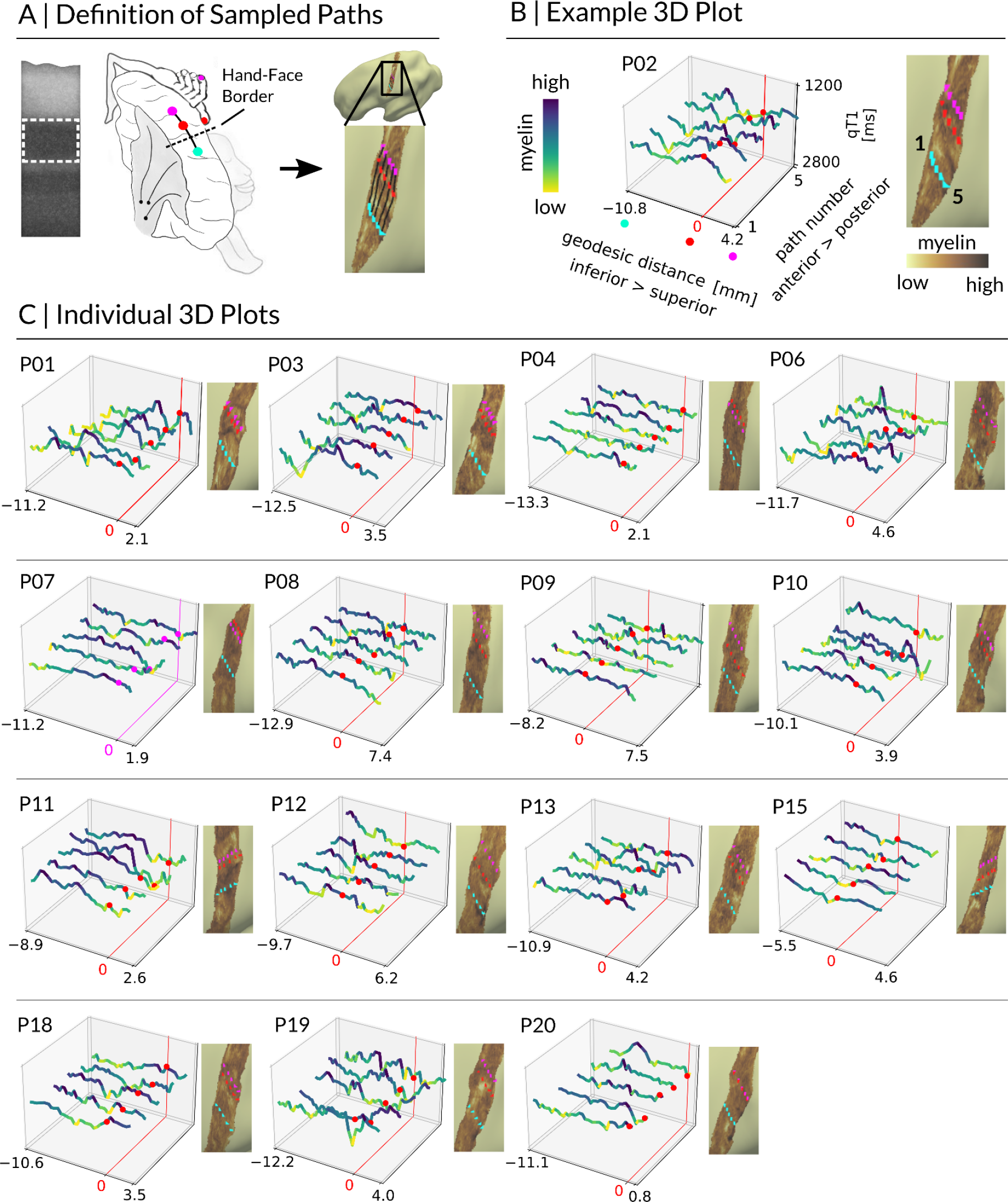
Multidimensional sampling of qT1-based myelin. (A) qT1 values sampled from the middle layer along multiple geodesic paths (black solid lines), connecting the upper face region (cyan) with D1 (red) and D2 (magenta). (B) Fluctuations in qT1-based myelin given in milliseconds (ms, z-axis) (yellow: low myelin, blue: high myelin) are shown in reference to geodesic distances given in millimeter (mm, x-axis) for multiple sampling locations (y-axis). Individual cortical surfaces show low myelin in lighter colors, high myelin in darker colors. The light-colored area between cyan and red dotted lines is also reflected by the yellow-colored qT1 “valleys” in the 3D plot. Red dots indicate the D1 representation (anchor to calculate geodesic distances). (C) Individual 3D plots of qT1 signals for n=15 participants.

In line with the results mentioned above, systematic low-myelin patterns (i.e. repetitive qT1 “valleys” which are visible in a minimum number of two neighboring sampling paths) were absent between D1 and D2 (see **Figures 8B, 8C**), except for 1/16 participants (P01), where this participant has been identified with stripe-like patterns between D1 and D2 in **Figure 6**. In two other cases (P11, P19), extracted D1 sampling points presumably crossed the hand-face border (**Figure 8C**), thus detected low-myelin patterns between D1 and D2 may actually reflect the hand-face border. In these cases, a shift of the low-myelin pattern from inferior to superior in reference to D1 (anchor to calculate geodesic distances) can be observed along the anterior-to posterior axis (i.e. across the sampled paths). We note that in one case (P20), where D1 and D2 sampling points almost overlapped (**Figure 8C**), the chance to detect a true border in the qT1 signal may have been reduced.

#### Non-Topographic 3D Structural Architecture of BA 3b Hand Area

To investigate whether finger representations differ in their 3D structural architecture, we compared layer-specific qT1, nQSM, pQSM and aQSM between individual finger representations using an ANOVA with the factors finger (D1, D2, D3, D4, D5) and layer (outer, middle, inner). For qT1 values, the analysis revealed a significant main effect of layer (F(1.24,23.51)=466.84, p=7.6⁻¹⁸, *·*^2^=0.82) with inner layers being more myelinated (showing lower qT1 values) than middle layers (inner: 1658.25 ± 6.35 (M ± SEM), middle: 1883.79 ± 7.13, t(19)=28.89, p=3.7⁻¹⁷, r=0.99), and middle layers being more myelinated than outer layers (outer: 2105.19 ± 11.79; t(19)=-14.58, p=9.0⁻¹², r=0.96; W=0, p=1.9⁻⁶), reflecting the intracortical myelin gradient described above. However, there was neither a significant main effect of finger (F(4,76)=1.54, p=0.2, *·*^2^=0.01) nor a significant interaction effect between layer and finger (F(3.92,74.53)=0.83, p=0.51, *·*^2^=0.004, see **Figure 9**).

**Figure 9.**
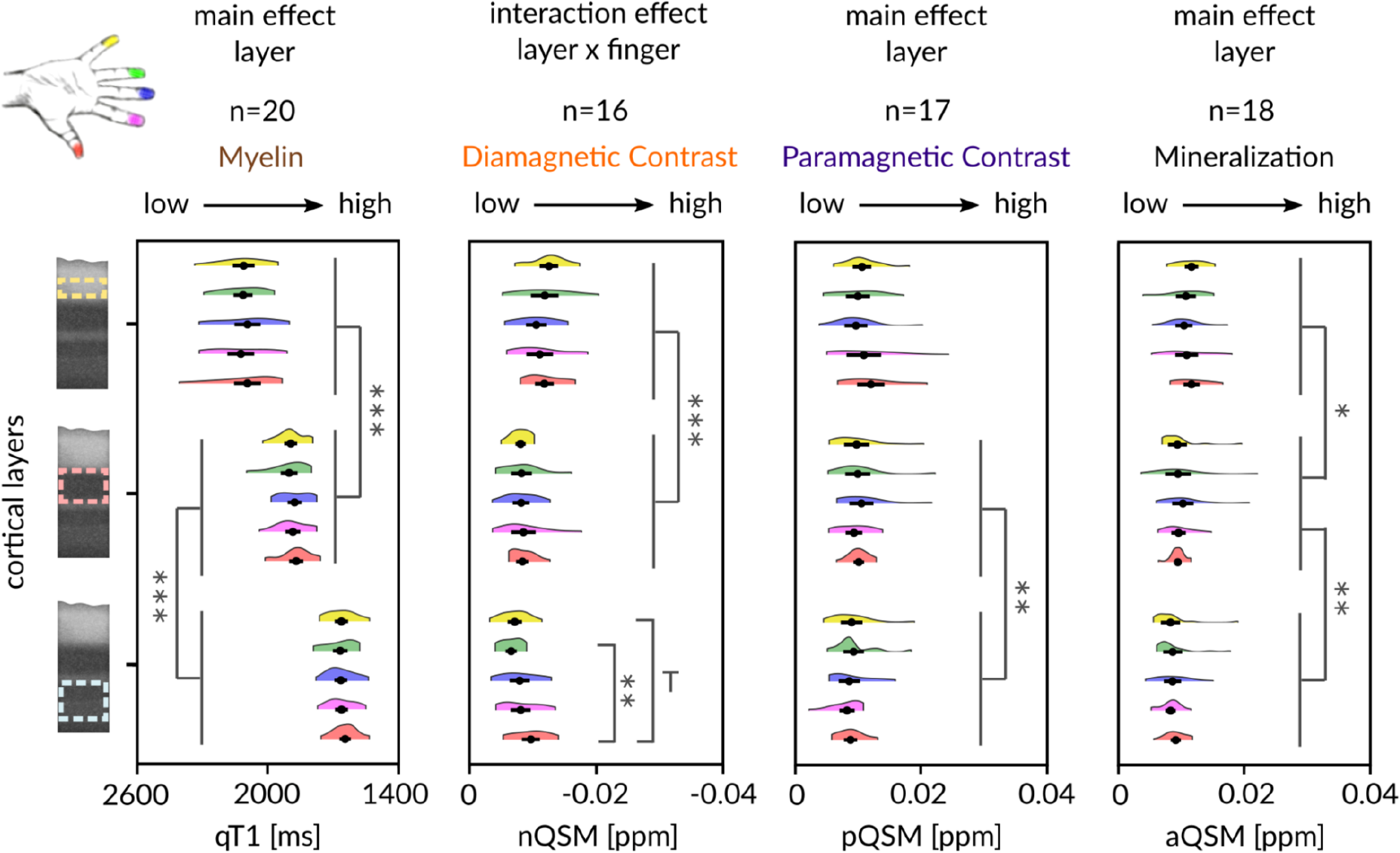
Non-topographic 3D structural architecture of the BA 3b hand area. Two-way repeated-measures ANOVAs (layer x finger) on qT1 (n=20), nQSM (n=16), pQSM (n=17) and aQSM (n=18) revealed significant differences in myelin content, diamagnetic contrast (e.g., calcium), paramagnetic contrast (iron) and mineralization content between layers (* p<0.025 , ** p<0.005, *** p<0.0005). Trends above Bonferroni-corrected threshold of p=0.005 correcting for 10 tests per layer to follow up a significant interaction are marked by a T. Fingers are shown in different colors: red: D1, magenta: D2, blue: D3, green: D4, yellow: D5. Except for higher nQSM values (higher diamagnetic contrast, second column) in D1 (red) compared to D4 (green) (significant difference with p=0.002) and D1 compared to D5 (yellow) (trend with p=0.007) in inner layers (light blue dotted line, bottom row), the 3D structural architecture with respect to myelin, paramagnetic contrast (iron) and mineralization does not significantly differ between the representations of single fingers in BA 3b. Myelin staining was remodeled according to Dinse et al. (2015) and shows anatomically-relevant layers (dotted lines, creme: outer layer, light pink: middle layer, light blue: inner layer) as reference. Black dots represent the mean, whiskers are drawn within the standard error of the mean.

The same ANOVA computed on pQSM values also showed a significant main effect of layer (F(1.28,20.51)=4.89, p=0.03, *·*^2^=0.05), here driven by middle layers (likely encompassing the input layer IV) showing higher pQSM values (more iron) than inner layers (likely encompassing layers V and VI; middle: 0.010 ± 5.8⁻⁴, inner: 0.009 ± 5.0⁻⁴; t(16)=3.84, p=0.001, r=0.69; W=14, p=0.002). Similar as for qT1 values, there was neither a significant main effect of finger (F(1.89,30.2)=0.32, p=0.71, *·*^2^=0.01) nor a significant interaction between layer and finger (F(2.81,44.88)=1.52, p=0.23, *·*^2^=0.02). The same ANOVA on aQSM values also showed a significant main effect of layer (F(1.46,24.86)=16.38, p=1.1⁻⁴, *·*^2^=0.12), here driven by outer layers showing more mineralization than middle layers (outer: 0.011 ± 4.2⁻⁴, middle: 0.010 ± 5.0⁻⁴; t(17)=3.09, p=0.007, r=0.60; W=22, p=0.004), and middle layers showing more mineralization than inner layers (inner: 0.009 ± 4.0⁻⁴; t(17)=3.71, p=0.002, r=0.67; W=19, p=0.002). However, similar as for qT1 and pQSM, there was neither a significant main effect of finger (F(2.44,41.54)=0.21, p=0.85, *·*^2^=0.004) nor a significant interaction effect (F(3.59,61.04)=1.07, p=0.38, *·*^2^=0.02).

Finally, the same ANOVA on nQSM values again showed a main effect of layer (F(2,30)=45.72, p=7.8⁻¹⁰, *·*^2^=0.26), here driven by more negative nQSM values (higher diamagnetic contrast) in outer compared to middle layers (outer: -0.012 ± 5.4⁻⁴, middle: -0.008 ± 4.7⁻⁴; t(15)=-9.37, p=1.2⁻⁷, r=0.92; W=0, p=3.1⁻⁵). Again, there was no significant main effect of finger (F(4,60)=0.97, p=0.43, *·*^2^=0.02), but a significant interaction between layer and finger (F(3.94,59.06)=2.78, p=0.04, *·*^2^=0.05). This interaction was driven by more negative nQSM values (higher diamagnetic tissue contrast) in the inner layers for D1 compared to D4 (D1: -0.010 ± 6.7⁻⁴, D4: -0.007 ± 4.2⁻⁴, t(15)=-3.82, p=0.002, r=0.70) and for D1 compared to D5 (D5: -0.007 ± 5.3⁻⁴, t(15)=-3.13, p=0.007, r=0.63).

In addition, we calculated the same ANOVAs after removing extreme outliers from pQSM and aQSM data. Again, the results show significant main effects of layer for pQSM values (F(1.44,18.66)=5.08, p=0.26, *·*^2^=0.07) and aQSM values (F(2,30)=29.47, p=8.3⁻⁸, *·*^2^=0.24).

We found higher pQSM values (higher iron content) in middle compared to inner layers (middle: 0.01 ± 4.7⁻⁴, inner: 0.008 ± 4.1⁻⁴, t(13)=3.37, p=0.005, r=0.68), higher aQSM values (higher mineralization) in outer compared to middle layers (outer: 0.011 ± 4.5⁻⁴, middle: 0.009 ± 3.4⁻⁴, t(15)=4.96, p=1.7⁻⁴, r=0.79) and in middle compared to inner layers (inner: 0.008 ± 2.4⁻⁴, t(15)=3.80, p=0.002, r=0.7). Again, there was neither a significant main effect of finger (pQSM: (F(2.41,31.35)=1.07, p=0.37, *·*^2^=0.03; aQSM: (F(4,60)=1.19, p=0.33, *·*^2^=0.02), nor a significant interaction effect between layer and finger (pQSM: (F(2.81,36.50)=1.68, p=0.19, *·*^2^=0.04; aQSM: (F(3.81,57.22)=1.53, p=0.21, *·*^2^=0.03).

Finally, we repeated the same ANOVAs for four equally-spaced layers (superficial, outer-middle, inner-middle, deep). Again, the results show significant main effects of layer for all tested substances (qT1: F(1.86,35.36)=581.16, p=2.4⁻²⁷, *·*^2^=0.80; nQSM: F(1.89,28.28)=21.03, p=3.53⁻⁶, *·*^2^=0.15; pQSM: F(1.98,31.72)=23.70, p=5.4⁻⁷, *·*^2^=0.12; aQSM: F(2.13,36.25)=5.16, p=.009, *·*^2^=0.03), but neither a significant main effect of finger (qT1: F(4,76)=1.85, p=0.13, *·*^2^=0.02; nQSM: F(2.60,39.06)=2.31, p=0.1, *·*^2^=0.05; pQSM: F(2.22,35.56)=0.41, p=0.69, *·*^2^=0.01; aQSM: F(2.19,37.24)=0.30, p=0.77, *·*^2^=0.01), nor a significant interaction between layer and finger (qT1: F(3.86,73.41)=1.06, p=0.38, *·*^2^=0.01; pQSM: F(4.79,76.61)=1.91, p=0.11, *·*^2^=0.02; aQSM: F(3.14,53.37)=0.59, p=0.63, *·*^2^=0.01), except for nQSM showing a trend for a significant interaction effect (nQSM: F(4.40,65.98)=2.34, p=0.058, *·*^2^=0.04).

Together, except for more diamagnetic tissue contrast in D1 compared to D4 and D5 in inner layers of BA 3b, our results show that the 3D structural architecture with respect to myelin, iron and mineralization does not significantly differ between the representations of single fingers in BA 3b.

#### High Similarity of Finger-Specific 3D Profiles

To investigate whether 3D structural profiles of different fingers are similar, we sampled qT1, nQSM, pQSM and aQSM values along 21 cortical depths specifically for each finger (see **Figure 10**). We compared intracortical meta parameters (i.e., skewness and kurtosis; in classical parcellation often used to differentiate between brain areas; e.g. Dinse et al., 2015) between finger representations. Using robust one-way repeated-measures ANOVAs on 20%-trimmed means with factor finger (D1, D2, D3, D4, D5), there were no significant differences in skewness and kurtosis of finger-specific 3D profiles (see **Figure 10B,** for exact statistical results see **Table 4**).

**Figure 10.**
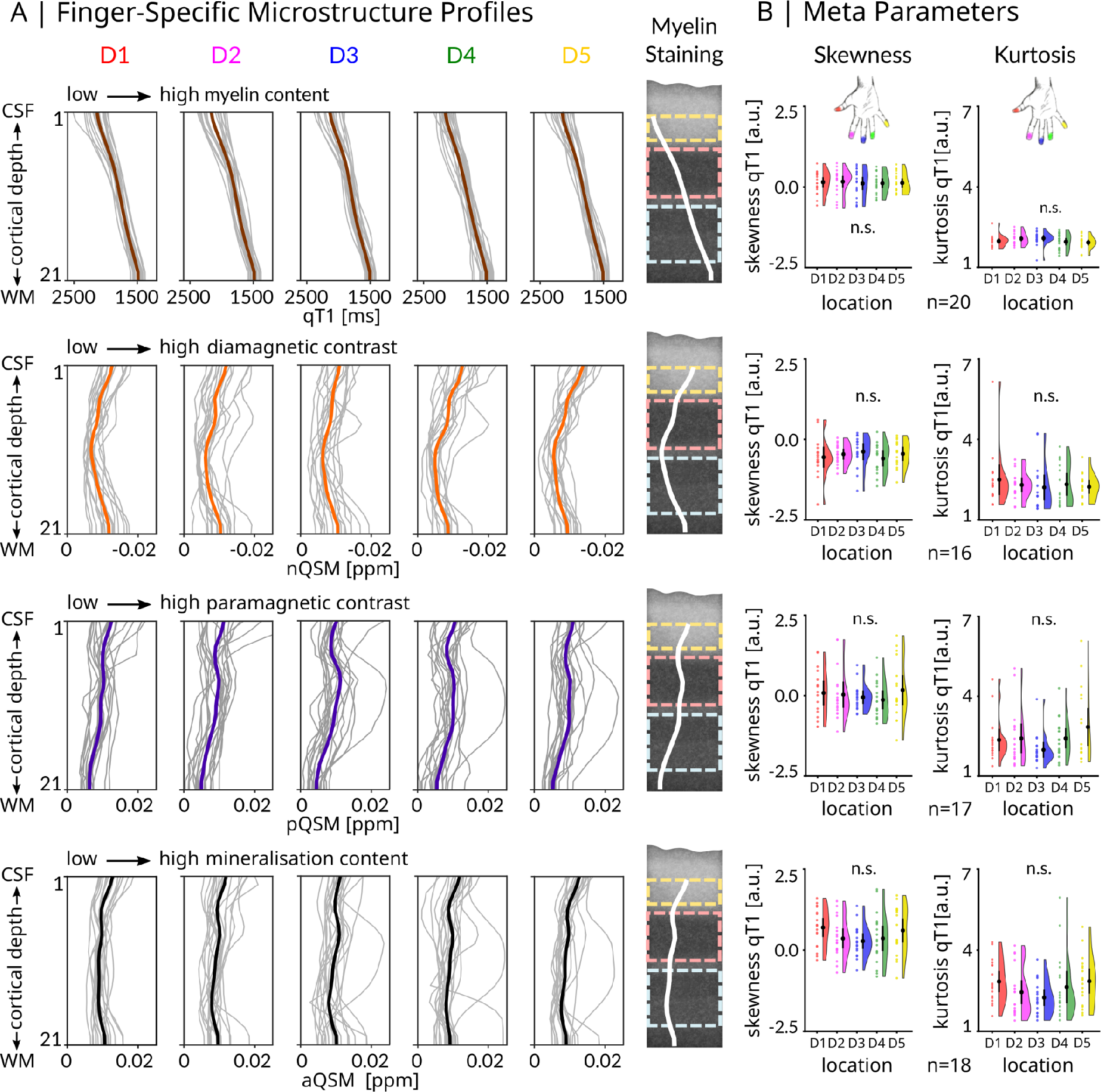
**Finger-specific 3D profiles and intracortical meta parameters do not differ between BA 3b finger representations**. (A) Quantitative T1 (qT1, reflecting myelin, n=20, group mean plotted in brown), negative QSM (nQSM, n=16, reflecting diamagnetic tissue contrast/calcium, group mean plotted in orange), positive QSM (pQSM, reflecting paramagnetic tissue contrast/iron, n=17, group mean plotted in purple) and absolute QSM (aQSM, reflecting mineralization, n=18, group mean plotted in black) sampled at 21 different cortical depths. Depth 1 is located closest to CSF, depth 21 is located closest to WM. Profiles of thumb (D1), index finger (D2), middle finger (D3), ring finger (D4), and little finger (D5) are shown. Myelin staining was remodeled according to Dinse et al. (2015) and shows anatomically-relevant cortical compartments (creme: outer layer, light pink: middle layer, light blue: inner layer) and average microstructure profiles across fingers (white lines) as reference. (B) Comparison of intracortical meta parameters between fingers using one-way repeated measures ANOVAs with within-subjects factor finger (red: D1, magenta: D2, blue: D3, green: D4, yellow: D5) revealed no significant (n.s.) differences in skewness and kurtosis between fingers. Values are given in arbitrary units (a.u.). Black dots represent the group mean. Whiskers (black lines) are drawn within the standard error of the mean. Colored dots represent individual data.

**Table 4.** Comparison of intracortical skewness and kurtosis values between fingers. There were no significant differences between fingers in intracortical skewness and kurtosis values (qT1, nQSM, pQSM, aQSM) revealed by one-way repeated-measures ANOVAs with within-subjects factor finger (D1, D2, D3, D4, D5), given normality (nQSM skewness, aQSM skewness). In all other cases (where data was non-normal), robust one-way repeated-measures ANOVAs on 20%-trimmed means are reported.

| MRI Proxy | Meta Parameter | n | DFn | DFd | F | p |
| --- | --- | --- | --- | --- | --- | --- |
| qT1 | skewness | 20 | 3.210 | 35.260 | 1.008 | 0.404 |
|  | kurtosis | 20 | 2.900 | 31.890 | 2.209 | 0.108 |
| nQSM | skewness | 16 | 4.000 | 60.000 | 0.571 | 0.685 |
|  | kurtosis | 16 | 2.940 | 26.480 | 0.483 | 0.693 |
| pQSM | skewness | 17 | 3.750 | 37.450 | 0.733 | 0.567 |
|  | kurtosis | 17 | 2.710 | 27.150 | 1.687 | 0.196 |
| aQSM | skewness | 18 | 4.000 | 68.000 | 1.427 | 0.234 |
|  | kurtosis | 18 | 4.000 | 44.000 | 1.474 | 0.226 |

More specifically, we calculated Fréchet distances for each participant between all possible pairs of finger-specific qT1 profiles as measure of profile similarity (see **Figure 11**). Similarity matrices were calculated both individually (see **Figure 11B**) and across participants (see **Figure 11C**). Overall, the results indicate high similarity between finger-specific 3D qT1 profiles. Note that values closer to zero reflect higher similarity. Using a robust one-way repeated measures ANOVA on 20%-trimmed means with factor finger pairs (levels: D1-D2, D2-D3, D3-D4, D4-D5), there were no significant differences in qT1 profile similarity between neighboring finger pairs (*F*(3,33)=0.32, p=0.81).

**Figure 11.**
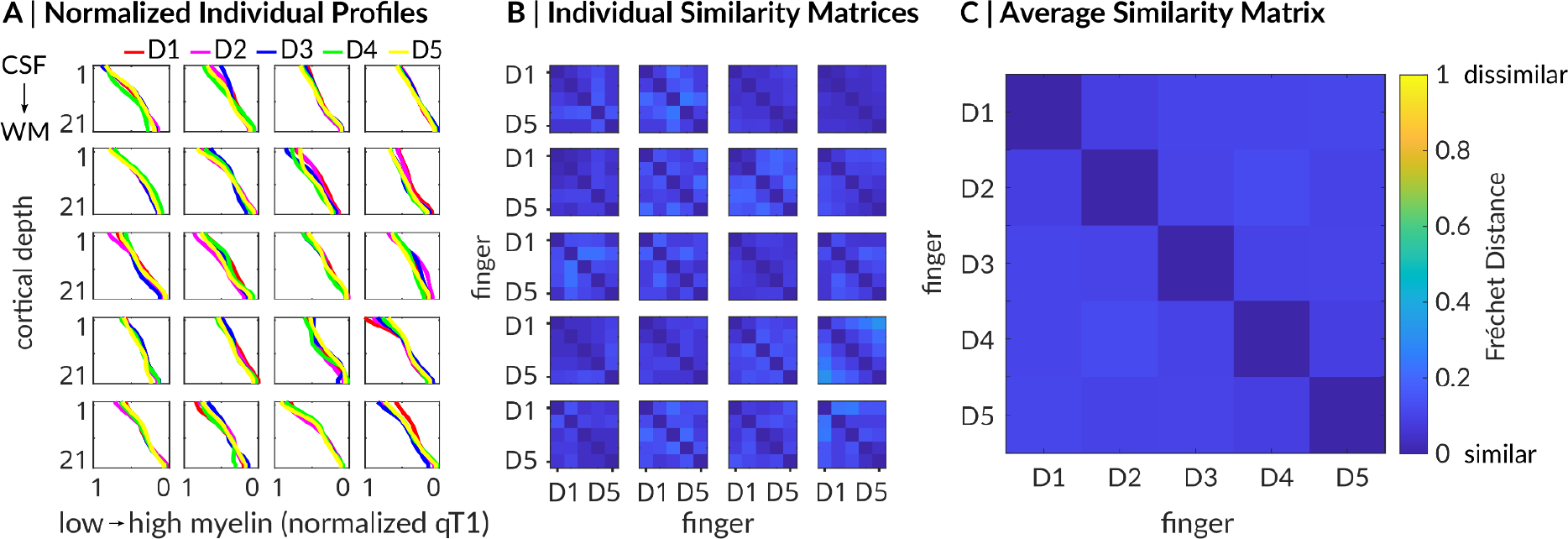
High similarity of finger-specific 3D qT1 profiles. (A) Individual finger-specific 3D profiles of normalized qT1 values (x-axis) sampled along 21 different cortical depths (y-axis). Different colors indicate different fingers (D1: thumb, D2: index finger, D3: middle finger, D4: ring finger, D5: little finger). Different panels reflect different participants (P01-P20). (B) Individual similarity matrices reflect individual Fréchet distances between finger profiles for all combinations of finger pairs (e.g., D1 vs. D2). Diagonals indicate perfect similarity (value of zero). Note that low values indicate high similarity (blue color). (C) Average similarity matrix of Fréchet distances (across participants) indicates high similarity of 3D profiles between individual fingers (blue color).

We also calculated Fréchet distances for each participant between all possible pairs of finger-specific nQSM, pQSM and aQSM profiles (see **Figure 12**). Overall, finger-specific QSM profiles appear visually less similar than finger-specific qT1 profiles. However, using robust one-way repeated measures ANOVAs on 20%-trimmed means with factor finger pairs (levels: D1-D2, D2-D3, D3-D4, D4-D5), we found no significant differences in similarity between neighboring fingers pairs, neither for nQSM profiles (F(3,27)=0.50, p=0.69), nor for pQSM profiles (F(1.77,17.67)=1.77, p=0.20), nor for aQSM profiles (F(2.85,31.34)=2.02, p=0.13).

**Figure 12.**
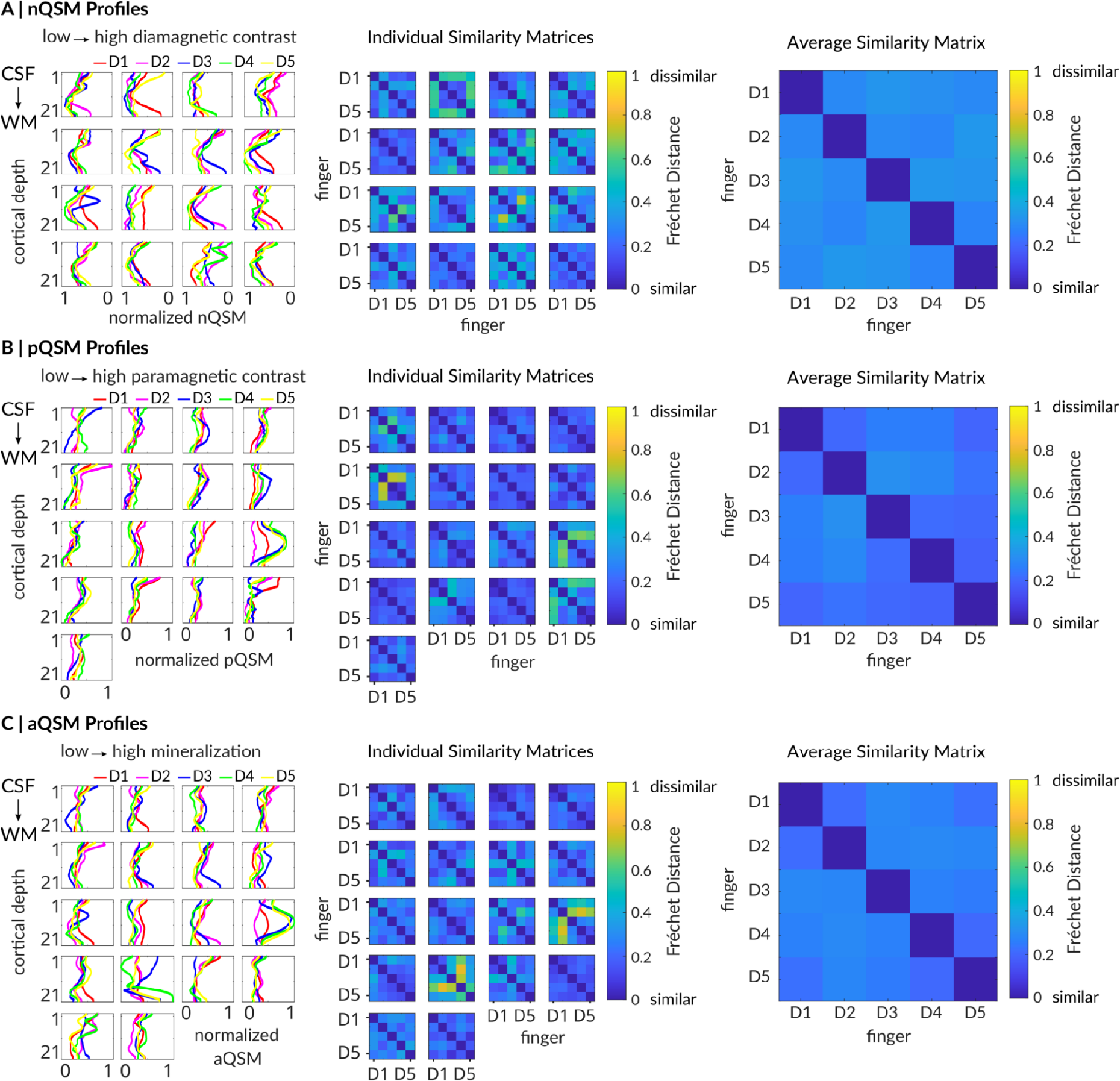
Similarity of finger-specific 3D QSM profiles. (A) Individual finger-specific 3D profiles of normalized negative QSM (nQSM) values (x-axis) sampled along 21 different cortical depths (y-axis). Different colors indicate different fingers (D1: thumb, D2: index finger, D3: middle finger, D4: ring finger, D5: little finger). Different panels reflect different participants. Individual similarity matrices reflect individual Fréchet distances between finger profiles (e.g., D1 vs. D2). Diagonals reflect perfect similarity (values of zero). Average similarity matrix of Fréchet distances (across participants) indicates similarity of nQSM profiles between individual fingers (blue color). (B) Individual finger-specific 3D profiles and similarity matrices of normalized positive QSM (pQSM) values. Average similarity matrix of Fréchet distances indicates similarity of pQSM profiles between individual fingers (blue color). (C) Individual finger-specific 3D profiles and similarity matrices of normalized absolute QSM (aQSM) values. Average similarity matrix of Fréchet distances indicates similarity of aQSM profiles between individual fingers (blue color).

Together, our results show that finger-specific 3D profiles of myelin, diamagnetic contrast, iron and mineralization do not significantly differ in their meta parameters between fingers.

#### Topographic 3D Structural Architecture of BA 3b Hand and Face Areas

To demonstrate that the methods used here are sensitive to capture topographic differences when they in fact exist, we carried out a control analysis comparing layer-specific qT1, pQSM, nQSM and aQSM between the functionally-localized face and hand areas (using action maps). A two-way repeated measures ANOVA on qT1 values with factors layer (inner, middle, outer) and body part (face, hand) revealed a significant interaction effect (F(1.37,20.55)=15.72, p=2.7⁻⁴, *·*^2^=0.02). qT1-based myelin content was lower in the hand compared to the face area in inner (face: 1601.19 ± 12.36, hand: 1638.62 ± 11.84, t(15)=2.91, p=0.011, r=0.6) and outer layers (face: 2067.11 ± 21.31, hand: 2097.74 ± 20.39, t(15)=2.22, p=0.042, r=0.5, see **Figure 13**). The same ANOVA computed on pQSM values revealed a significant main effect of body part (F(1,13)=10.92, p=0.006, *·*^2^=0.12). pQSM-based iron content was highest in the hand area (hand: 0.011 ± 4.3⁻⁴, face: 0.009 ± 3.6⁻⁴). For aQSM values there was also a significant main effect of body part (F(1,13)=10.92, p=0.006, *·*^2^=0.13), showing highest mineralization content in the hand area (hand: 0.010 ± 3.5⁻⁴, face: 0.009 ± 3.3⁻⁴); whereas for nQSM values there was no significant difference between body parts.

**Figure 13.**
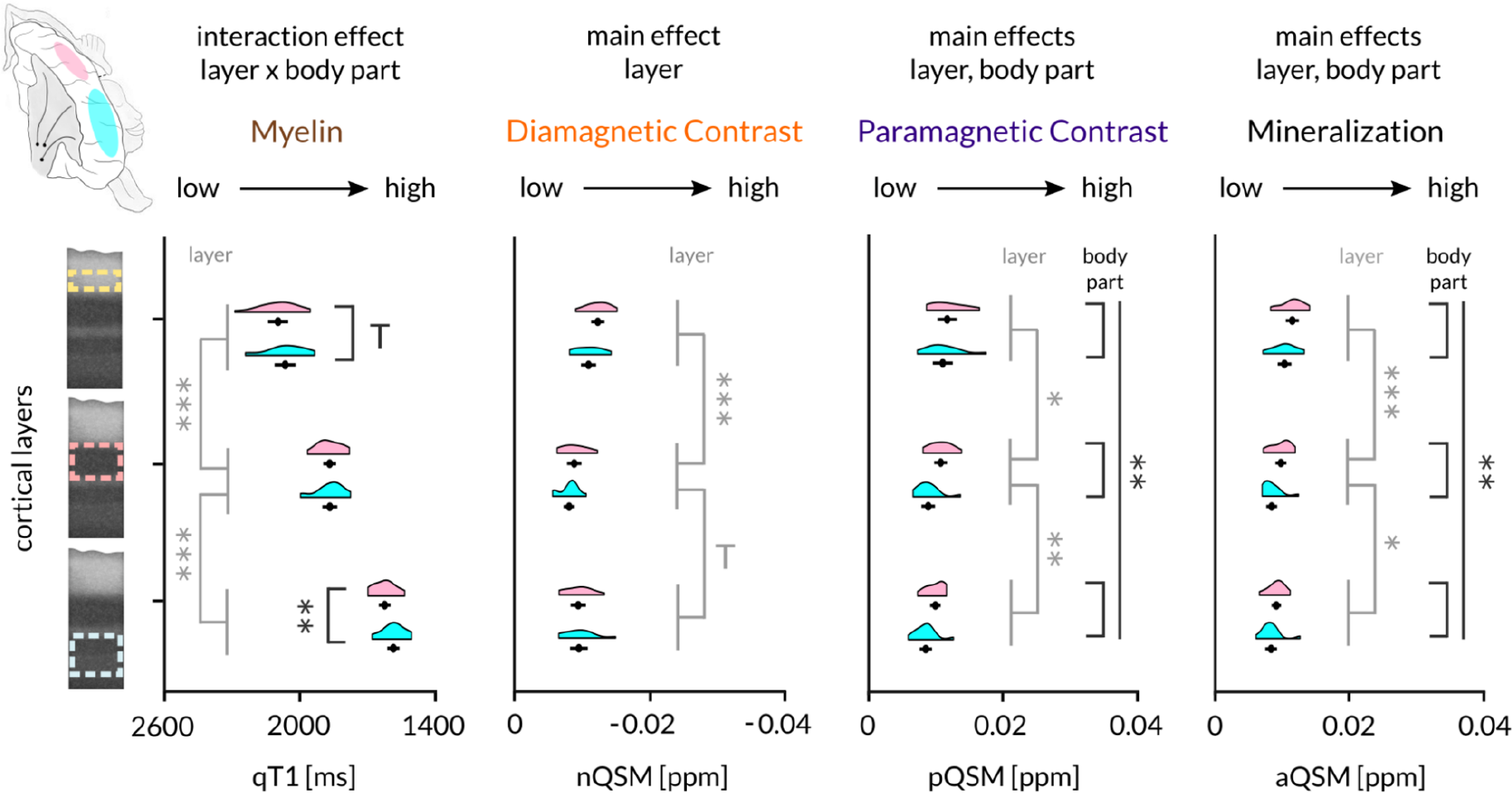
Topographic 3D structural architecture of the BA 3b face-hand area. wo-way repeated-measures ANOVAs (layer x body part) on qT1 (n=16), nQSM (n=14), pQSM (n=14) and aQSM (n=14) revealed significant differences in myelin content, diamagnetic contrast (e.g., calcium), paramagnetic contrast (iron) and mineralization content between layers (significance statement colored in light gray, p < 0.025 *, p < 0.017 **, p < 0.001 ***). Trends above Bonferroni-corrected threshold of p=0.017 correcting for 3 tests per layer to follow up a significant interaction are marked by a T. Body parts are shown in different colors (light pink: hand, cyan: face). Except for nQSM (diamagnetic contrast), the 3D structural architecture, with respect to myelin, paramagnetic contrast (iron) and mineralization, differs significantly between the representations of the hand and face in BA 3b (significance statement colored in dark gray). Myelin staining was remodeled according to Dinse et al. (2015) and shows anatomically-relevant layers (dotted lines, creme: outer layer, light pink: middle layer, light blue: inner layer). Black dots represent the mean, whiskers are drawn within the standard error of the mean.

Together, our results show that the 3D structural architecture, with respect to myelin, iron and mineralization, significantly differs between major body part representations (here the representations of the face and the hand in BA 3b) whereas, as explained above, such differences were not detected between individual fingers.

## Discussion

We used sub-millimeter quantitative MRI proxies for myelin, iron and mineralization to answer the question of whether the 3D architecture of the human BA 3b hand area is homogenous, or whether there are systematic microstructural differences between finger representations. We show that precise intra-cortical tissue contrasts can be extracted from a group of younger adults and can be used to define anatomically-relevant layer compartments. While low-myelin borders were visible between the representations of the hand and the face, such borders were not systematically detected between finger representations. Similarly, while the 3D structural architecture significantly differed between the representations of the hand and the face, it did not differ significantly between finger representations (except for deep layers of the thumb) or along the inferior-to-superior axis. In fact, we found 3D structural profiles to be highly similar between individual finger representations. Taken together, our data show that the 3D structural architecture of the human BA 3b hand area is homogenous in younger adults and essentially non-topographic, although functional finger representations are distinct and discontinuous. This difference compared to the structural architecture of the hand area in some monkey species, and to the hand-face area in humans, reveals novel insights into the neuronal mechanisms that underlie the architecture and flexibility of finger representations and cortical plasticity in humans.

As expected from histology (e.g. Vogt and Vogt, 1919) and *in vivo ex vivo* validation work (Stüber et al., 2014; Dinse et al., 2015), we show an intra-cortical myelin gradient with increasing myelin content (decreasing qT1 values) from superficial to deep cortical depth. By relating our data to remodeled *in vivo ex vivo* validated data (Dinse et al., 2015), we extracted three anatomically-relevant cortical compartments that contained estimates for BA 3b layers IV and V/VI. This approach is novel, because in previous work, data of only few participants were investigated extensively (e.g., Alkemade et al., 2022; Huber et al., 2020), or layers were defined arbitrarily by dividing the cortex into equally-spaced compartments (Kuehn et al., 2017; Tardiff et al., 2015). While our layer definition was based on cortical myelination (i.e., qT1 values), the pQSM and nQSM data also fit to this compartmentalization scheme, since the two iron peaks coincide with the expected locations of the Baillarger bands in anatomical layers IV and V, and the u-shaped nQSM profile shows a small plateau where we expected layer IV to be located. This scheme was therefore used to describe the 3D architecture of BA 3b in more detail.

With respect to low-myelin borders, we do not find systematic evidence for such borders between finger representations in BA 3b, which seems to differentiate humans from some monkey species (Jain, Catania and Kaas, 1998; Qi and Kaas, 2004). While topographic borders have been related to reducing plasticity (Sereno, 2005), an absence of structural borders may introduce flexibility to finger (re-)alignment. In older humans’ BA 3b, more overlap between neighboring finger representations and reduced cortical distances between index and middle finger representations have been described (Liu et al., 2021). In older rats, however, upper limb and whisker representations that are separated by septa remain intact (Spengler et al., 1995; Coq and Xerri, 2000; Godde et al., 2002; David-Jürgens et al., 2008). In reelin-deficient mice, where the primary visual cortex is intra-cortically disorganized, visual cortical plasticity is enhanced (Pielecka-Fortuna et al., 2015). Thus, an absence of sharp structural borders (both vertically and horizontally) may allow more distributed information (Muret et al., 2022), facilitating cortical flexibility.

An absence of low-myelin borders between BA 3b finger representations has also conceptual implications. Although single fingers are functionally distinct units, our findings suggest that all fingers are represented as one structural unit. Low-myelin borders in the cortex may therefore separate cortical representations that are nearby in the cortex but distant in the real word, as previously shown for the hand and face (Glasser et al., 2016; Kuehn et al., 2017; Northall et al., 2023). When cortical representations are nearby in the cortex and also nearby in the real word, such as for individual fingers, GABA-ergic functional inhibition may allow their differentiation when they receive distinct input (Kuehn et al., 2014). Low-myelin borders may therefore reflect real world features. This also explains why structural and functional parcellation does not always align (Zhi et al., 2002), and relates to the concept that local stability and global reorganization of finger representations are driven by distributed, rather than finger-specific, processing underlying the topographic map (Wesselink et al., 2022). An absence of low-myelin borders between BA 3b finger representations is therefore in line with recent observations, and suggests that the hand area is encoded as an integrated unit, shaped in close relation to real world features such as external spatial location (Haggard et al., 2006).

In human M1, large-scale body part representations are not only separated by low-myelin borders, but also differ in their microstructural profiles (Northall et al., 2023). In BA 3b, both were present between the hand and the face, but absent between individual fingers (except for the thumb). This indicates that low-myelin borders and 3D structural differences coincide (or are both absent), adding evidence that large-scale body part representations can be regarded as distinct cortical fields (Sereno et al., 2022).

The exception to the rule in our data is the thumb, which shows higher diamagnetic contrast in deeper layers compared to the ring and the little finger. Most neurons in the BA 3b thumb representation specifically respond to tactile stimulation of the thumb, but not to tactile stimulation of other fingers, and interconnect only sparsely to other digit representations. Also in macaque monkeys, the thumb representation may act as an independent module processing selective information, whereas information content is more distributed between other fingers (Lazar et al., 2023). This may be linked to its special role for tool use.

Previous research in humans demonstrated that brain areas of similar cortical myelo- and cytoarchitecture are more likely to be connected and tend to exhibit stronger cortico-cortical connections (Huntenburg et al., 2017; Wei et al., 2018). The high similarity of finger-specific 3D structural profiles emphasizes cortical spread between finger representations, which may facilitate changes in finger topography, for example after synchronous stimulation or gluing of multiple fingers (Kalisch et al. 2008, Kolasinski et al. 2016), and may offer a mechanistic explanation as to why finger representations are unstable in older age (Liu et al. 2021; Kuehn et al. 2018b). Our findings encourage the investigation of microstructure profiles to fully understand topographic plasticity in aging and neurodegenerative diseases (Schreiber et al. 2021).

Addressing study limitations, absence of evidence should not be equated with evidence of absence. It still remains possible that potential low-myelin borders between finger representations could not be detected with the given methodology. To overcome possible confounds, additional descriptive and multidimensional analyses were implemented. Future studies may use pattern recognition algorithms to achieve fully automated septa-like feature detection. A direct methodological validation using the hand-face septum was not possible due to a lack of forehead activation during our motor task (Root et al. 2022). Future studies may consider isolated movements of the eyebrow to detect the hand-face septum in BA 3b. Although the sample size of this study is high compared to previous 7T-MRI studies in humans and histological studies in monkey species on the topic (e.g. Kuehn et al., 2017; Huber et al., 2020; Qi and Kaas, 2004; Jain, Catania and Kaas, 1998), it is relatively low for group statistics. Consequently, type I and type II errors are more difficult to control (Sullivan et al., 2016). To reduce type I error, we corrected for multiple comparisons and restricted the number of statistical tests to the minimum number needed. To reduce type II errors, we show statistical trends (i.e., p > 0.05 and < 0.1, see Figures 4, 9, 13) (Liu et al. 2021), and one-sided tests were applied to directional hypotheses (i.e. low-myelin borders). To increase sensitivity for the null hypothesis (no effect), we conducted Bayesian statistics. However, it still remains possible that vertices of BA 1 and/or BA 3a were included in the statistics, even though we carefully checked segmentation results.

Taken together, our data provide the first comprehensive *in vivo* description of the 3D structural architecture of the human BA 3b hand area and show it to be non-topographic. This distinguishes the human hand area from the “barrel field” of rodents and the hand area of monkeys, which show low-myelin borders separating individual whiskers and fingers, respectively. The specific homogeneous structural organization of the human hand area in BA 3b suggests that there are less structural limitations to cortical plasticity and reorganization. Studying the individual microstructure of the cortex together with functional changes is critical to fully understand the mechanisms of topographic change in the course of aging, learning, or disease, and may also explain individual differences in these mechanisms. These new insights encourage future studies to incorporate the dimension of cortical layers into new testable models, and consider structural limitations of plasticity for intervention methods.

## Acknowledgments

This project was funded by the German Research Foundation (Deutsche Forschungsgemeinschaft, DFG) (KU 3711/2-1, project number: 423633679 and Project-ID 425899996 – SFB 1436). A.N. was funded by the Else Kröner Fresenius Stiftung: 2019-A03. We thank Lilith-Sophie Lange and Miriam Weber for their support in data collection, Pierre-Louis Bazin and Daniel Haenelt for giving advice on ultra high-resolution image processing, Susanne Stoll for discussions about multilevel modeling and Nikolaus Kinder for graphical support.

## References

1. Acosta-Cabronero J, Betts MJ, Cardenas-Blanco A, Yang S, Nestor PJ (2016) *In Vivo* MRI Mapping of Brain Iron Deposition across the Adult Lifespan. The Journal of Neuroscience 36:364–374.

2. Acosta-Cabronero J, Milovic C, Mattern H, Tejos C, Speck O, Callaghan MF (2018) A robust multi-scale approach to quantitative susceptibility mapping. Neuroimage 183:7–24.

3. Alkemade A, Bazin P-L, Balesar R, Pine K, Kirilina E, Möller HE, Trampel R, Kros JM, Keuken MC, Bleys RLAW, Swaab DF, Herrler A, Weiskopf N, Forstmann BU (2022) A unified 3D map of microscopic architecture and MRI of the human brain. Sci Adv 8:eabj7892.

4. Alt H, Godau M (1995) Computing the Fréchet distance between two polygonal curves. Int J Comput Geom Appl 05:75–91.

5. Avants BB, Tustison NJ, Song G, Cook PA, Klein A, Gee JC (2011) A reproducible evaluation of ANTs similarity metric performance in brain image registration. Neuroimage 54:2033–2044.

6. Bazin P-L, Pham DL (2007) Topology-preserving tissue classification of magnetic resonance brain images. IEEE Trans Med Imaging 26:487–496.

7. Bazin PL, Weiss M, Dinse J, Schäfer A, Trampel R, Turner R (2014) A computational framework for ultra-high resolution cortical segmentation at 7Tesla. Neuroimage 93 Pt 2:201--209.

8. Blake DT, Byl NN, Merzenich MM (2002) Representation of the hand in the cerebral cortex. Behav Brain Res 135:179–184.

9. Bürkner PC (2018) Advanced Bayesian Multilevel Modeling with the R Package brms. The R Journal 10:395.

10. Bürkner PC (2017) brms: An R Package for Bayesian Multilevel Models Using Stan. J Stat Softw 80:1–28.

11. Carpenter B, Gelman A, Hoffman MD, Lee D, Goodrich B, Betancourt M, Brubaker MA, Guo J, Li P, Riddell A (2017) Stan: A Probabilistic Programming Language. J Stat Softw 76.

12. Cohen J (1988) Statistical Power Analysis for the Behavioral Sciences, 2nd Edition. Routledge.

13. Coq JO, Xerri C (2000) Age-related alteration of the forepaw representation in the rat primary somatosensory cortex. Neuroscience 99:403–411.

14. David-Jürgens M, Churs L, Berkefeld T, Zepka RF, Dinse HR (2008) Differential effects of aging on fore- and hindpaw maps of rat somatosensory cortex. PLoS One 3:e3399.

15. Dinse J, Härtwich N, Waehnert MD, Tardif CL, Schäfer A, Geyer S, Preim B, Turner R, Bazin P-L (2015) A cytoarchitecture-driven myelin model reveals area-specific signatures in human primary and secondary areas using ultra-high resolution in-vivo brain MRI. Neuroimage 114:71–87.

16. Donatelli G, Costagli M, Cecchi P, Migaleddu G, Bianchi F, Frumento P, Siciliano G, Cosottini M (2022) Motor cortical patterns of upper motor neuron pathology in amyotrophic lateral sclerosis: A 3 T MRI study with iron-sensitive sequences. Neuroimage Clin 35:103138.

17. Elbert T, Pantev C, Wienbruch C, Rockstroh B, Taub E (1995) Increased cortical representation of the fingers of the left hand in string players. Science 270:305–307.

18. Feldman DE, Brecht M (2005) Map plasticity in somatosensory cortex. Science 310:810–815.

19. Field A, Miles J, Field Z (2012) Discovering statistics using R. London, England: SAGE Publications.

20. Florence SL, Jain N, Kaas JH (1997) Plasticity of Somatosensory Cortex in Primates. Seminars in Neuroscience 9:3–12.

21. Geyer S, Schleicher A, Zilles K (1999) Areas 3a, 3b, and 1 of human primary somatosensory cortex. Neuroimage 10:63–83.

22. Glasser MF, Coalson TS, Robinson EC, Hacker CD, Harwell J, Yacoub E, Ugurbil K, Andersson J, Beckmann CF, Jenkinson M, Smith SM, Van Essen DC (2016) A multi-modal parcellation of human cerebral cortex. Nature 536:171–178.

23. Glasser MF, Van Essen DC (2011). Mapping human cortical areas in vivo based on myelin content as revealed by T1- and T2-weighted MRI. J Neurosci 31:11597–11616.

24. Godde B, Berkefeld T, David-Jürgens M, Dinse HR (2002) Age-related changes in primary somatosensory cortex of rats: evidence for parallel degenerative and plastic-adaptive processes. Neurosci Biobehav Rev 26:743–752.

25. Haacke EM, Xu Y, Cheng Y-CN, Reichenbach JR (2004) Susceptibility weighted imaging (SWI). Magn Reson Med 52:612–618.

26. Haggard P, Kitadono K, Press C, Taylor-Clarke M (2006) The brain’s fingers and hands. Exp Brain Res 172:94–102.

27. Hametner S, Endmayr V, Deistung A, Palmrich P, Prihoda M, Haimburger E, Menard C, Feng X, Haider T, Leisser M, Köck U, Kaider A, Höftberger R, Robinson S, Reichenbach JR, Lassmann H, Traxler H, Trattnig S, Grabner G (2018) The influence of brain iron and myelin on magnetic susceptibility and effective transverse relaxation - A biochemical and histological validation study. Neuroimage 179:117–133.

28. Han X, Pham DL, Tosun D, Rettmann ME, Xu C, Prince JL (2004) CRUISE: cortical reconstruction using implicit surface evolution. Neuroimage 23:997–1012.

29. Huber L, Finn ES, Handwerker DA, Bönstrup M, Glen DR, Kashyap S, Ivanov D, Petridou N, Marrett S, Goense J, Poser BA, Bandettini PA (2020) Sub-millimeter fMRI reveals multiple topographical digit representations that form action maps in human motor cortex. Neuroimage 208:116463.

30. Huntenburg JM, Bazin P-L, Goulas A, Tardif CL, Villringer A, Margulies DS (2017) A Systematic Relationship Between Functional Connectivity and Intracortical Myelin in the Human Cerebral Cortex. Cereb Cortex 27:981–997.

31. Huntenburg JM, Bazin P-L, Margulies DS (2018) Large-Scale Gradients in Human Cortical Organization. Trends Cogn Sci 22:21–31.

32. In M-H, Posnansky O, Speck O (2016) PSF mapping-based correction of eddy-current-induced distortions in diffusion-weighted echo-planar imaging. Magn Reson Med 75:2055–2063.

33. Jain N, Catania KC, Kaas JH (1998) A histologically visible representation of the fingers and palm in primate area 3b and its immutability following long-term deafferentations. Cereb Cortex 8:227–236.

34. Kalisch T, Tegenthoff M, Dinse HR (2008) Improvement of sensorimotor functions in old age by passive sensory stimulation. Clin Interv Aging 3:673–690.

35. Kolasinski J, Makin TR, Jbabdi S, Clare S, Stagg CJ, Johansen-Berg H (2016) Investigating the Stability of Fine-Grain Digit Somatotopy in Individual Human Participants. J Neurosci 36:1113–1127.

36. Kuehn E, Dinse J, Jakobsen E, Long X, Schäfer A, Bazin P-L, Villringer A, Sereno MI, Margulies DS (2017) Body Topography Parcellates Human Sensory and Motor Cortex. Cereb Cortex 27:3790–3805.

37. Kuehn E, Haggard P, Villringer A, Pleger B, Sereno MI (2018a) Visually-Driven Maps in Area 3b. J Neurosci 38:1295–1310.

38. Kuehn E, Mueller K, Turner R, Schütz-Bosbach S (2014) The functional architecture of S1 during touch observation described with 7 T fMRI. Brain Struct Funct 219:119–140.

39. Kuehn E, Perez-Lopez MB, Diersch N, Döhler J, Wolbers T, Riemer M (2018b) Embodiment in the aging mind. Neurosci Biobehav Rev 86:207–225.

40. Kuehn E, Pleger B (2020) Encoding schemes in somatosensation: From micro- to meta-topography. Neuroimage 223:117255.

41. Langkammer C, Schweser F, Krebs N, Deistung A, Goessler W, Scheurer E, Sommer K, Reishofer G, Yen K, Fazekas F, Ropele S, Reichenbach JR (2012) Quantitative susceptibility mapping (QSM) as a means to measure brain iron? A post mortem validation study. Neuroimage 62:1593–1599.

42. Lazar L, Chand P, Rajan R, Mohammed H, Jain N (2023) Somatosensory cortex of macaque monkeys is designed for opposable thumb. Cereb Cortex 33:195–206.

43. Liu P, Chrysidou A, Doehler J, Hebart MN, Wolbers T, Kuehn E (2021) The organizational principles of de-differentiated topographic maps in somatosensory cortex. Elife 10:e60090.

44. Lucas BC, Bogovic JA, Carass A, Bazin P-L, Prince JL, Pham DL, Landman BA (2010) The Java Image Science Toolkit (JIST) for rapid prototyping and publishing of neuroimaging software. Neuroinformatics 8:5–17.

45. Mair P, Wilcox R (2020) Robust statistical methods in R using the WRS2 package. Behav Res Methods 52:464–488.

46. Marques JP, Kober T, Krueger G, van der Zwaag W, Van de Moortele P-F, Gruetter R (2010) MP2RAGE, a self bias-field corrected sequence for improved segmentation and T1-mapping at high field. Neuroimage 49:1271–1281.

47. Martuzzi R, van der Zwaag W, Farthouat J, Gruetter R, Blanke O (2014) Human finger somatotopy in areas 3b, 1, and 2: a 7T fMRI study using a natural stimulus. Hum Brain Mapp 35:213–226.

48. McAuliffe MJ, Lalonde FM, McGarry D, Gandler W, Csaky K, Trus BL (2001) Medical Image Processing, Analysis and Visualization in clinical research. In: Proceedings 14th IEEE Symposium on Computer-Based Medical Systems. CBMS 2001, pp 381–386.

49. Meyer HS, Egger R, Guest JM, Foerster R, Reissl S, Oberlaender M (2013) Cellular organization of cortical barrel columns is whisker-specific. Proc Natl Acad Sci U S A 110:19113–19118.

50. Muret D, Root V, Kieliba P, Clode D, Makin TR (2022) Beyond body maps: Information content of specific body parts is distributed across the somatosensory homunculus. Cell Rep 38:110523.

51. Nasreddine ZS, Phillips NA, Bédirian V, Charbonneau S, Whitehead V, Collin I, Cummings JL, Chertkow H (2005) The Montreal Cognitive Assessment, MoCA: a brief screening tool for mild cognitive impairment. J Am Geriatr Soc 53:695–699.

52. Northall A, Doehler J, Weber M, Vielhaber S, Schreiber S, Kuehn E (2023) Layer-Specific Vulnerability is a Mechanism of Topographic Map Aging. bioRxiv 2022.05.29.493865.

53. Oldfield RC (1971) The assessment and analysis of handedness: the Edinburgh inventory. Neuropsychologia 9:97–113.

54. Paquola C, Vos De Wael R, Wagstyl K, Bethlehem RAI, Hong S-J, Seidlitz J, Bullmore ET, Evans AC, Misic B, Margulies DS, Smallwood J, Bernhardt BC (2019) Microstructural and functional gradients are increasingly dissociated in transmodal cortices. PLoS Biol 17:e3000284.

55. Peters A (2009) The effects of normal aging on myelinated nerve fibers in monkey central nervous system. Front Neuroanat 3:11.

56. Petersen CCH (2007) The functional organization of the barrel cortex. Neuron 56:339–355.

57. Petersen CCH, Crochet S (2013) Synaptic computation and sensory processing in neocortical layer 2/3. Neuron 78:28–48.

58. Pielecka-Fortuna J, Wagener RJ, Martens A-K, Goetze B, Schmidt K-F, Staiger JF, Löwel S (2015) The disorganized visual cortex in reelin-deficient mice is functional and allows for enhanced plasticity. Brain Struct Funct 220:3449–3467.

59. Pinheiro J, Bates D, R Core Team (2022) nlme: Linear and Nonlinear Mixed Effects Models. Available at: https://CRAN.R-project.org/package=nlme.

60. Pleger B, Wilimzig C, Nicolas V, Kalisch T, Ragert P, Tegenthoff M, Dinse HR (2016) A complementary role of intracortical inhibition in age-related tactile degradation and its remodelling in humans. Sci Rep 6:27388.

61. Puckett AM, Bollmann S, Junday K, Barth M, Cunnington R (2020) Bayesian population receptive field modeling in human somatosensory cortex. Neuroimage 208:116465.

62. Qi H-X, Kaas JH (2004) Myelin stains reveal an anatomical framework for the representation of the digits in somatosensory area 3b of macaque monkeys. J Comp Neurol 477:172–187.

63. R Core Team (2022) R: A Language and Environment for Statistical Computing. Available at: https://www.R-project.org/.

64. Ragert P, Schmidt A, Altenmüller E, Dinse HR (2004) Superior tactile performance and learning in professional pianists: evidence for meta-plasticity in musicians. European Journal of Neuroscience 19:473–478.

65. Root V, Muret D, Arribas M, Amoruso E, Thornton J, Tarall-Jozwiak A, Tracey I, Makin TR (2022) Complex pattern of facial remapping in somatosensory cortex following congenital but not acquired hand loss. eLife 11:e76158.

66. Schellekens W, Thio M, Badde S, Winawer J, Ramsey N, Petridou N (2021) A touch of hierarchy: population receptive fields reveal fingertip integration in Brodmann areas in human primary somatosensory cortex. Brain Struct Funct 226:2099–2112.

67. Schreiber S, Northall A, Weber M, Vielhaber S, Kuehn E (2021) Topographical layer imaging as a tool to track neurodegenerative disease spread in M1. Nat Rev Neurosci 22:68–69.

68. Schweisfurth MA, Frahm J, Schweizer R (2015) Individual left-hand and right-hand intra-digit representations in human primary somatosensory cortex. Eur J Neurosci 42:2155–2163.

69. Schweisfurth MA, Schweizer R, Frahm J (2011) Functional MRI indicates consistent intra-digit topographic maps in the little but not the index finger within the human primary somatosensory cortex. Neuroimage 56:2138–2143.

70. Schweisfurth MA, Schweizer R, Treue S (2014) Feature-based attentional modulation of orientation perception in somatosensation. Front Hum Neurosci 8:519.

71. Schweizer R, Voit D, Frahm J (2008) Finger representations in human primary somatosensory cortex as revealed by high-resolution functional MRI of tactile stimulation. Neuroimage 42:28–35.

72. Schwenkreis P, El Tom S, Ragert P, Pleger B, Tegenthoff M, Dinse HR (2007) Assessment of sensorimotor cortical representation asymmetries and motor skills in violin players. Eur J Neurosci 26:3291–3302.

73. Sereno MI (2005) Neuroscience: plasticity and its limits. Nature 435:288–289.

74. Sereno MI, Sood MR, Huang R-S (2022) Topological Maps and Brain Computations From Low to High. Front Syst Neurosci 16:787737.

75. Sethian JA (1999) Level Set Methods and Fast Marching Methods: Evolving Interfaces in Computational Geometry, Fluid Mechanics, Computer Vision, and Materials Science. Cambridge University Press.

76. Shoham D, Grinvald A (2001) The cortical representation of the hand in macaque and human area S-I: high resolution optical imaging. J Neurosci 21:6820–6835.

77. Spengler F, Godde B, Dinse HR (1995) Effects of ageing on topographic organization of somatosensory cortex. Neuroreport 6:469–473.

78. Stringer EA, Qiao P-G, Friedman RM, Holroyd L, Newton AT, Gore JC, Min Chen L (2014) Distinct fine-scale fMRI activation patterns of contra- and ipsilateral somatosensory areas 3b and 1 in humans. Hum Brain Mapp 35:4841–4857.

79. Stüber C, Morawski M, Schäfer A, Labadie C, Wähnert M, Leuze C, Streicher M, Barapatre N, Reimann K, Geyer S, Spemann D, Turner R (2014) Myelin and iron concentration in the human brain: a quantitative study of MRI contrast. Neuroimage 93:95–106.

80. Sullivan C, Kaszynski A (2019) PyVista: 3D plotting and mesh analysis through a streamlined interface for the Visualization Toolkit (VTK). J Open Source Softw 4:1450.

81. Sullivan LM, Weinberg J, Keaney JF Jr (2016) Common Statistical Pitfalls in Basic Science Research. J Am Heart Assoc 5.

82. Tardif CL, Schäfer A, Waehnert M, Dinse J, Turner R, Bazin P-L (2015) Multi-contrast multi-scale surface registration for improved alignment of cortical areas. Neuroimage 111:107–122.

83. Tosun D, Rettmann ME, Han X, Tao X, Xu C, Resnick SM, Pham DL, Prince JL (2004) Cortical surface segmentation and mapping. Neuroimage 23 Suppl 1:S108–S118.

84. Ursell T (2022) Frechet Distance Calculator. Available at: https://de.mathworks.com/matlabcentral/fileexchange/41956-frechet-distance-calculator [Accessed August 30, 2022].

85. Valk SL, Xu T, Paquola C, Park B-Y, Bethlehem RAI, Vos de Wael R, Royer J, Masouleh SK, Bayrak S, Kochunov P, Yeo BTT, Margulies D, Smallwood J, Eickhoff SB, Bernhardt BC (2022) Genetic and phylogenetic uncoupling of structure and function in human transmodal cortex. Nat Commun 13:2341.

86. Vogt C, Vogt O (1919) Allgemeine ergebnisse unserer hirnforschung. JA Barth.

87. Waehnert MD, Dinse J, Weiss M, Streicher MN, Waehnert P, Geyer S, Turner R, Bazin P-L (2014) Anatomically motivated modeling of cortical laminae. Neuroimage 93 Pt 2:210–220.

88. Wagstyl K, Ronan L, Goodyer IM, Fletcher PC (2015) Cortical thickness gradients in structural hierarchies. Neuroimage 111:241–250.

89. Wei Y, Scholtens LH, Turk E, van den Heuvel MP (2019) Multiscale examination of cytoarchitectonic similarity and human brain connectivity. Netw Neurosci 3:124–137.

90. Welker C (1976) Receptive fields of barrels in the somatosensory neocortex of the rat. J Comp Neurol 166:173–189.

91. Welker C, Woolsey TA (1974) Structure of layer IV in the somatosensory neocortex of the rat: description and comparison with the mouse. J Comp Neurol 158:437–453.

92. Wesselink DB, Sanders Z-B, Edmondson LR, Dempsey-Jones H, Kieliba P, Kikkert S, Themistocleous AC, Emir U, Diedrichsen J, Saal HP, Makin TR (2022) Malleability of the cortical hand map following a finger nerve block. Sci Adv 8:eabk2393.

93. Woolsey TA, Van der Loos H (1970) The structural organization of layer IV in the somatosensory region (SI) of mouse cerebral cortex. The description of a cortical field composed of discrete cytoarchitectonic units. Brain Res 17:205–242.

94. Yousry TA, Schmid UD, Alkadhi H, Schmidt D, Peraud A, Buettner A, Winkler P (1997) Localization of the motor hand area to a knob on the precentral gyrus. A new landmark. Brain 120 Pt 1:141–157.

95. Zeidman P, Silson EH, Schwarzkopf DS, Baker CI, Penny W (2018) Bayesian population receptive field modelling. Neuroimage 180:173–187.

96. Zheng W, Nichol H, Liu S, Cheng Y-CN, Haacke EM (2013) Measuring iron in the brain using quantitative susceptibility mapping and X-ray fluorescence imaging. Neuroimage 78:68–74.

97. Zhi D, King M, Hernandez-Castillo CR, Diedrichsen J (2022) Evaluating brain parcellations using the distance-controlled boundary coefficient. Hum Brain Mapp 43:3706–3720.

